# Senescent stromal cells promote cancer resistance through SIRT1 loss-potentiated overproduction of small extracellular vesicles

**DOI:** 10.1101/2020.03.22.002667

**Authors:** Liu Han, Qilai Long, Shenjun Li, Qixia Xu, Boyi Zhang, Xuefeng Dou, Min Qian, Yannasittha Jiramongkol, Jianming Guo, Liu Cao, Y. Eugene Chin, Eric W-F Lam, Jing Jiang, Yu Sun

## Abstract

Cellular senescence is a potent tumor-suppressive program that prevents neoplastic events. Paradoxically, senescent cells develop an inflammatory secretome, termed the senescence-associated secretory phenotype (SASP) and implicated in age-related pathologies including cancer. Here we report that senescent cells actively synthesize and release small extracellular vesicles (sEVs) with a distinctive size distribution. Mechanistically, SIRT1 loss supports accelerated sEV production despite enhanced proteome-wide ubiquitination, a process correlated with ATP6V1A downregulation and defective lysosomal acidification. Once released, senescent stromal sEVs significantly alter the expression profile of recipient cancer cells and enhance their aggressiveness, specifically drug resistance mediated by expression of ATP binding cassette subfamily B member 4 (ABCB4). Targeting SIRT1 with an agonist SRT2104 prevents development of cancer resistance through restraining sEV production by senescent stromal cells. In clinical oncology, sEVs in peripheral blood of posttreatment cancer patients are readily detectable by routine biotechniques, presenting a novel biomarker to monitor therapeutic efficacy and to predict long term outcome. Together, our study identifies a distinct mechanism supporting pathological activities of senescent cells, and provides a novel avenue to circumvent advanced human malignancies by co-targeting cancer cells and their surrounding microenvironment, which contributes to drug resistance via secretion of sEVs from senescent stromal cells.

## INTRODUCTION

Cytotoxic chemotherapy represents an effective modality for cancer treatment, inducing clinical responses associated with a significantly lowered risk of recurrence, but only in a limited fraction of patients ^1–3^. As selection of genes accurately predicting therapeutic responses might improve cancer outcomes, gene signatures aimed at predicting responses to specific anticancer agents and minimizing drug resistance are being actively evaluated ^4^. Multiple studies have reported side-effects generated by local and systemic treatments, and their therapeutic benefits may be restrained by tumor-promoting host responses induced by certain forms of cytotoxicity ^5^. Development of innate and/or acquired resistance are among the major mechanisms underlying diminished responsiveness to anticancer regimens, as disease relapse after an initial response can occur as a result of systemic release of multiple soluble factors, which frequently remodel the treatment-damaged tumor microenvironment (TME) ^6, 7^.

Cellular senescent is a state of cell cycle arrest that occurs upon exposure of cells to different stresses, usually resistant to cell death induction. While cellular senescence is beneficial for a few physiological events such as tissue repair, wound healing and embryogenesis, it accelerates organismal aging and is responsible for age-related disorders ^8^. Senescent cells synthesize and secrete a plethora of extracellular proteins, covering growth factors, cytokines, chemokines and proteases, a phenomenon referred to as the SASP ^9–11^. Expression of most of the SASP components are regulated by kinases including TAK1, p38, mTOR and Jak2/Stat3, all of which indeed converge on the NF-κB complex and the c-EBP/β family of transcription factors ^12^. By means of the SASP expression, senescent cells remodel the local tissue via paracrine mechanisms and recruit immune cells in aging tissues ^13^. In response to damage signals, degradation of the nuclear lamina and elimination of the major structural component lamin B1 not only enhance histone depletion and chromatin remodeling but also forms microtubule-associated protein light chain 3 (LC3/ATG8)-containing cytoplasmic chromatin fragments ^14, 15^. If not eliminated via exosome secretion, such chromatin fragments can activate the cyclic GMP–AMP synthase (cGAS) and stimulator of interferon-γ (IFN-γ) genes (STING) response, further promoting expression of the SASP and type I interferons (IFNs) ^16–19^.

Extracellular vesicles (EVs) are membranous vesicles released by almost all cell types and contain a lipid bilayer that protects the luminal contents of proteins, nucleic acids, lipids and metabolites against harsh environmental conditions ^20^. According to their distinctive biogenesis and release modes, EVs can be categorized as exosomes, microvesicles (MVs) and apoptotic bodies (ABs) ^21^. The cargos packaged within or associated with the EVs can reflect the pathophysiological state of host cells ^22^. However, the capacity of senescent cell-derived EVs in orchestrating intercellular communications within the TME and changing pathological track under therapeutic settings, remain largely unexplored. To this end, we investigated the biogenesis mechanisms of EVs generated by senescent human stromal cells developing the SASP, and disclosed that EVs can shape acquired resistance of cancer cells to anticancer treatments. We further demonstrated the feasibility of modulating EV production to control therapeutic resistance acquired from the treatment-damaged TME, thus presenting a novel avenue to improve therapeutic efficacy by harnessing senescent cell-derived EVs.

## RESULTS

### Senescent stromal cells generate an increased number of EVs with distinct size distribution

As stromal cells represent the major non-cancerous mesenchymal components providing structural architecture within the tissue of most solid tumors, we chose to employ a primary normal human prostate stromal cell line, namely PSC27, to initiate the study. Composed of predominantly fibroblasts but with a minor percentage of non-fibroblast cell lineages including endothelial cells and smooth muscle cells, PSC27 develops a typical SASP after exposure to stressful insults such as cytotoxic chemotherapy or ionizing radiation ^23–25^. We treated these cells with a pre-optimized sub-lethal dose of bleomycin (BLEO), resulting in enhanced senescence-associated β-galactosidase (SA-β-Gal) staining positivity, decreased BrdU incorporation, and elevated DNA damage foci several days afterwards (Fig. S1A-C). Data from Illumina sequencing (RNA-seq) indicated that PSC27 cells were strongly expressing multiple SASP factors including but not limited to CXCL8, CCL20, CSF3, IL-1α and CCL3 after treatment (Fig. S1D). We isolated stromal cell-derived vesicles with sequential ultracentrifugation before assessment of their size and distribution. Strikingly, quantitative nanoparticle tracking analysis (NTA) showed that the number of senescent (SEN) PSC27 cell-released vesicles between 30 nm and 1.0 μm, a size range characterized of major eukaryotic EV subtypes including exosomes and MVs ^26^, was approximately 7 times of their pre-senescent (PRE) counterparts (Fig. 1A). Further, we noticed a substantial difference of EV size distribution between PRE and SEN stromal cells. In contrast to EVs from PRE cells, which were mainly enriched at 72, 102, 121 and 170 nm, EVs from SEN cells displayed an evident re-distribution in particle diameter, as reflected by several major peaks at 33, 47, 64, 122, 156, 182, 266 and 314 nm, respectively, although in each case the refereed EVs fell in the category of small EVs (sEV) as defined by recent literatures ^26, 27^ (Fig. 1B). The average size shifted from 116 nm to 161 nm, with a statistical significance between PRE- and SEN-derived sEVs (Fig. 1C-D). Although there was a small number of EVs displaying sizes beyond such a principal window, they accounted for a limited percentage of overall stromal EVs and were not the main focus of subsequent investigations.

**Fig. 1.**
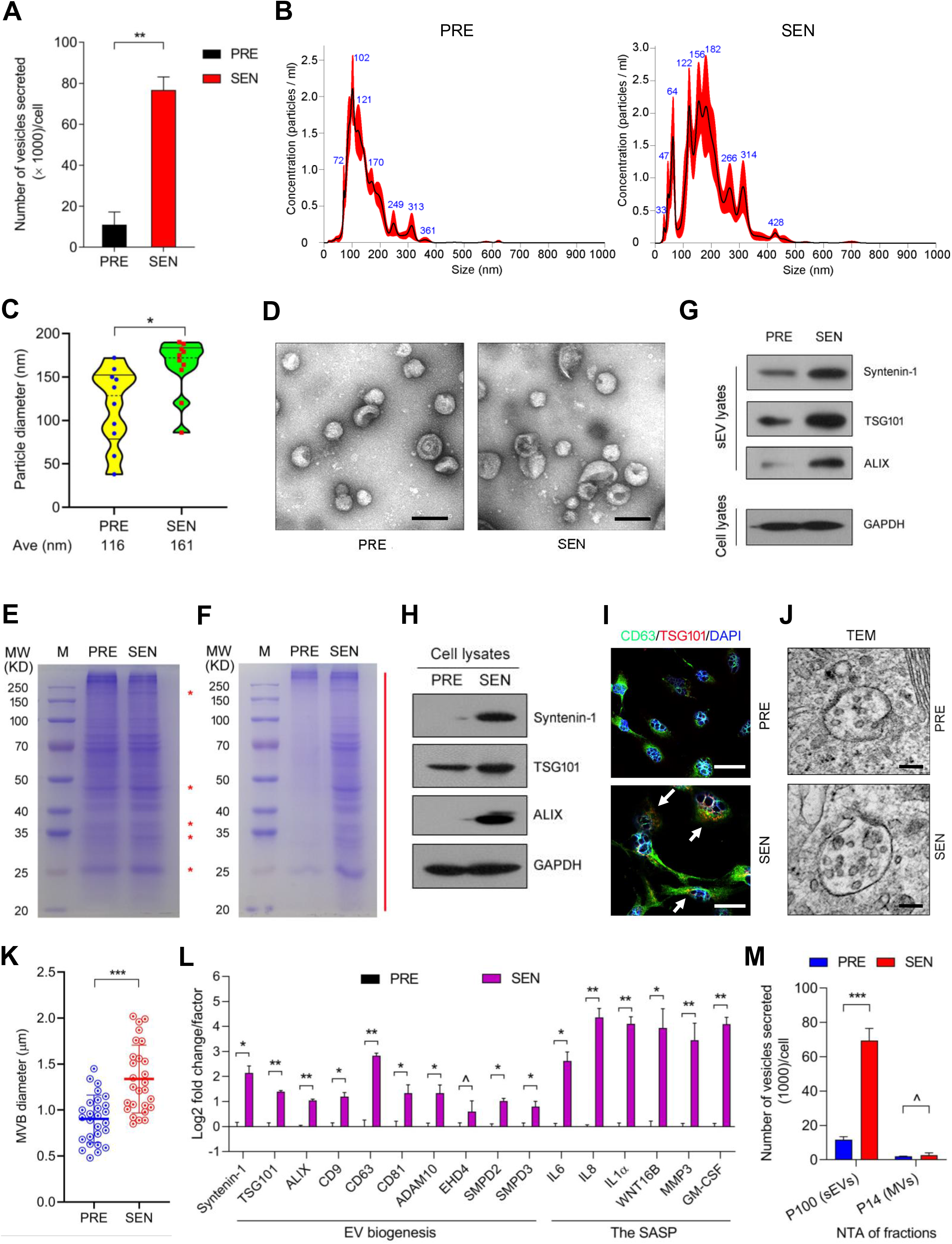
Stromal cells produce an increased number of sEVs with distinct size distribution upon senescence. (**A**) Quantitative comparison of the number of sEVs secreted from PRE and SEN PSC27 cells in 3 consecutive days and detected by NTA. Data normalized to cell number. (**B**) Concentration (mean ± SD; n = 5 acquisitions of one sample per condition) and size distribution of sEVs calculated by NTA. Left, PRE; Right, SEN. (**C**) Average diameter of sEVs secreted by PRE and SEN cells. Data from NTA assessment. (**D**) Representative transmission electron microscopy (TEM) images of sEVs isolated from stromal cells (n = 3 independent biological samples). Scale bars, 100 nm. (**E**) Coomassie brilliant blue stained SDS-PAGE gel which shows visible difference at multiple sizes (red stars) between PRE and SEN stromal cell sEVs. Lysates loaded upon normalization to the number of sEVs. (**F**) Coomassie brilliant blue stained protein gel, with sEV lysates loaded upon normalization to the number of parental cells. Red line, the range of proteins that show remarkable difference. (**G**) Immunoblot analysis of stromal cell-derived total sEVs collected in 3 days, with the lysate loading normalized to parental cell number. (**H**) Immunoblot analysis of whole lysates of parental cells. (**I**) Immunofluorescence staining of PRE and SEN stromal cells with CD63 and TSG101. Scale bars, 10 μm. Arrows, sEVs in synthesis (TSG101 positive). (**J**) TEM of stromal cells, with the representative images showing multivesicular bodies (MVBs) in PRE and SEN cells, respectively. Scale bars, 300 nm. (**K**) Comparative statistics of the diameter of MVBs in PRE vs SEN stromal cells, with MVBs representative of each cell state selected for measurement. (**L**) Quantitative expression assay of sEV biomarker and biogenesis-related molecules, with SASP hallmark factors assessed in parallel. (**M**) NTA measurement of the number of P100-EVs (sEVs) and P14-EVs (MVs) released from PRE and SEN stromal cells, after successive differential ultracentrifugation, with the data normalized to cell number per vehicle subtype. ashok, *P* > 0.05; *, *P* < 0.05; **, *P* < 0.01; ***, *P* < 0.001.

Preliminary in-gel analysis indicated that protein components of sEVs from PRE and SEN cells apparently differ from each other, when sample loading was normalized to the number of sEVs (Fig. 1E). However, when loading was normalized to the number of parental cells that secreted these sEVs, we noticed substantially enhanced amounts of sEV protein cargoes (Fig. 1F). The findings were supported by immunoblots, which suggested that expression of sEV-specific markers such as syntenin-1, TSG101 and ALIX was markedly increased in total sEV lysates upon cellular senescence (Fig. 1G). We then analyzed the whole lysates of parental cells, and found considerably elevated expression of sEV markers in SEN cells, in contrast to PRE samples (Fig. 1H). Upon immunofluorescence (IF) staining, we found the expression levels of CD63 (a tetraspanin protein) and TSG101 apparently enhanced in SEN cells, implying the possibility of active synthesis of sEVs in MVBs (Fig. 1I). Electron microscopy imaging showed expanding MVBs with an increase in both volume and the number of their internal vesicles in SEN cells, relative to their proliferating counterparts (Fig. 1J-K). Indeed, there was a concomitant upregulation of the vast majority of sEV-associated molecules as indicated by transcript assays, although the expression level of SMPD2/3, two neutral sphingomyelinases (nSMases) associated with biogenesis of both sEVs and MVs, appeared increased, as well (Fig. 1L). Further analysis with successive differential ultracentrifugation of pellets from conditioned media, a strategy that effectively separates sEVs from MVs (14,000 g and 100,000 g of sedimentation, respectively) ^28^, indicated enhanced secretion of sEVs by SEN cells, while MV release remained largely unchanged, implying engagement of a mechanism specifically responsible for production of sEVs, but not all EV subtypes (Fig. 1M). Notably, appearance of such a distinct expression pattern was accompanied by pronounced upregulation of several hallmark SASP factors including but not limited to IL6, IL8, IL1α, WNT16B, MMP3 and GM-CSF (Fig. 1L).

To validate the data, we chose HBF1203, another stromal cell line which was isolated from human breast tissue and consists mainly of fibroblasts. We treated HBF1203 with doxorubicin (DOX), a chemotherapeutic agent frequently used in breast cancer clinics. The results suggested that senescence-associated changes including those in cell phenotypes, sEV marker and SASP factor expression, and sEV production, can be readily reproduced in these cells (Fig. S1E-J). Thus, these data uncover a common feature potentially shared by stromal cells of multiple tissue and/or organ origin.

### Senescent stromal sEVs exhibit significantly altered spectrum and expression of small RNA species

EVs carry abundant bioactive components, including multiple types of small RNAs (sRNAs) specifically microRNAs (miRNAs), which can image the pathophysiological state of parental cells and frequently participate in intercellular communications ^29, 30^. We speculated that sEVs isolated from SEN stromal cells may differ substantially from their proliferating counterparts. To prove this hypothesis, we performed small RNA sequencing (sRNA-seq) to define the transcriptomic profile of sRNA species in stromal cell-derived sEVs. The data suggested that the total number of sRNAs essentially did not change. However, the number of certain sRNA species showed remarkable alterations, such as those falling in the categories of rRNA or tRNA, and those with genomic loci mapped in the intron of human genes (Fig. 2A; Table S1).

**Fig. 2.**
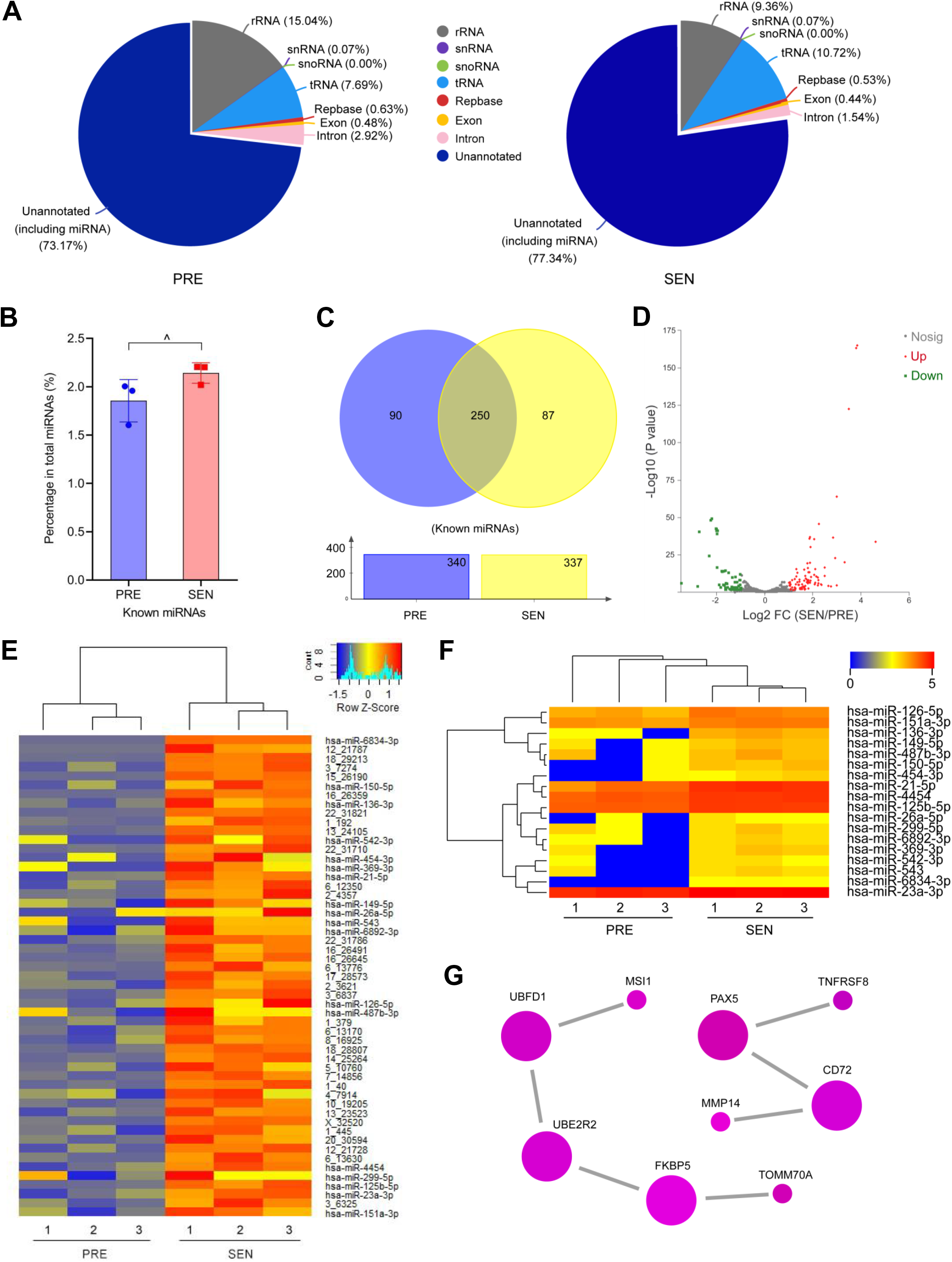
Transcriptome-wide expression profiling of small RNAs carried by senescent stromal cell-derived sEVs. (**A**) Comparative statistics of small RNA species in sEVs of PSC27 cells as detected by sRNA-seq. Left, PRE cells; right, SEN cells. (**B**) Statistical presentation of the percentage of known miRNAs, which have published sequences and annotations searchable in miRBase (http://www.mirbase.org/), among all sRNAs sequenced per cell population. (**C**) Venn diagram showing the known miRNAs carried by sEVs from PRE and SEN stromal cells, with the total miRNA number per cell population and the number of miRNA overlapping between two populations presented. (**D**) Volcano plot presenting the downregulated and upregulated miRNAs (totally 170) in SEN stromal cells. Green, downregulated; red, upregulated; grey, insignificantly changed. (**E**) Heatmap depicting the top 53 miRNAs (both known or novel) that were significantly upregulated in SEN vs PRE stromal cells, with an average fold change > 2.0 (*P* < 0.01 by *t* test). (**F**) Heatmap and hierarchical clustering showing the top 18 known miRNAs significantly upregulated in SEN vs PRE stromal cells (*P* < 0.01 by Mann-Whitney *U* test). (**G**) Protein-protein interaction network of the central molecular targets of the 18 most upregulated known miRNAs. Data from calculation and filtering by miRanda, TargetScan and RNAhybrid databases.

We next focused on miRNAs, a unique sRNA subspecies that can be delivered via sEVs to impact the fate, function and behavior of recipient cells ^31, 32^. Between PRE and SEN cells, there was an overlap of 834 miRNAs (Fig. S2A). Among the 1031 miRNAs presented by Illumina data, we found that the percentage of hitherto characterized miRNA (hereafter known miRNAs) in the total miRNA pool remained largely consistent, between sEVs derived from PRE and SEN cells (Fig. 2B). Between PRE and SEN cells, there were 250 known miRNAs in the overlap zone (Fig. 2C). We then focused on the 170 miRNAs that were significantly up- or down-regulated in SEN stromal cells (53 up and 117 down, respectively (Fig. 2D and Fig. S2B). Of note, there was a relatively consistent panel of most abundantly expressed miRNAs in either PRE or SEN cell populations, regardless of their annotated identity (known vs novel) (Fig. S2C-D).

Among the 53 miRNAs significantly upregulated in SEN cells, the relevance of neither the known or novel transcripts in cellular senescence has been extensively documented by available literatures (Fig. 2E). Although the 18 miRNA transcripts on the top list have been characterized by various biological implications and functional activities, they appeared clustered together with close proximity as revealed by hierarchical appraisal (Fig. 2F). We performed miRNA target prediction with miRanda and TargetScan programs and found 28 target proteins co-presented by NR, Swiss-Prot, EggNOG, GO, KEGG and Pfam databases which together cover 290 molecules in the overlapping zone (Fig. S2E). GO functional analysis suggest that molecular targets of these known miRNAs carried by SEN stromal sEVs are generally involved in regulation of biological, cellular and metabolic processes (Fig. S2F). KEGG database mining suggest that many of these targets are associated with mannose biosynthesis, TNF, AMPK and MAPK signaling pathways (Fig. S2G).

We further noticed that miRNAs of SEN stromal sEVs hold the potential to target proteins that actively participate in cell apoptosis. For example, hsa-miR-6834-3p (targeting THAP1, NAIF1, PERP, PDCD1 and BID), has-miR-150-5p (targeting AIFM2 and PDCD4), hsa-miR-136-3p (targeting CASP7/8/10, PDCD4/5 and FAS) and has-miR-542-3p (targeting CDIP1 and DEDD2) (Table S2). However, upon global assessment of protein-protein interaction (PPI), we found the presence of a mutually interactive network composed mainly of TNFRSF8, CD72, TOMM70A, MSI, UBE2R2, PAX5, FKBP5, MMP14, UBFD1 (Fig. 2G; Table S3). Many of these molecules are associated with apoptosis induction (TNFRSF8), intracellular beta-catenin degradation (UBE2R2) and inhibition of mTOR signaling pathway (FKBP5), processes largely of anticancer relevance. Thus, miRNAs delivered by SEN stromal sEVs may have the capacity to enhance recipient cell survival and circumvent cell death in intracellularly stressful and/or environmentally adverse conditions.

### SIRT1 decline supports deficient lysosomal acidification and enhanced sEV biogenesis via ATP6V1A downregulation

Given the substantial change of the number and cargo composition of sEVs generated by stromal cells upon senescence, we interrogated the mechanism supporting these alterations. It was suggested that dysfunctional lysosome or autophagy can promote EV biogenesis through modifying the fate of MVBs, the direct cytoplasmic source of exosomes ^33, 34^. A recent study further revealed that reduced SIRT1 expression in breast cancer cells can modify lysosomal activity, resulting in enhanced release of exosomes from these cells and significant changes in their exosome composition ^35^.

We first assessed the expression level of human SIRT family, a group of NAD^+^-dependent deacetylases, in proliferating and senescent cells. Whole transcriptomic analysis by RNA-seq suggested that all 7 members of this family were ubiquitously downregulated, although a statistical significance was observed mainly in SIRT1 and SIRT2 (Fig. 3A-B). Downregulation of SIRT1/2 was accompanied by upregulation of IL8 and MMP3, hallmark SASP factors (Fig. 3C). Upon cellular senescence, DNA damage-associated concomitant decline of SIRT1/2 was indeed also observed in a human primary fibroblast line BJ ^36^, essentially supporting our data. SIRT1 regulates diverse cellular targets and correlates with organismal aging, while SIRT2 has been reported to be marker of cellular senescence in certain cancer types such as osteosarcoma ^37, 38^. Although loss of SIRT1 enhances the secretion of exosomes by breast cancer cells, whether or not this occurs in stromal cells remains unclear. In our work, knockdown of SIRT1 with small hairpin RNAs (shRNAs) caused significant upregulation of not only the typical sEV markers including TSG101, Syntenin-1, ALIX and tetraspenins (CD9/63/81), but also downregulation of HSP70, the latter reported to be responsible for accumulation of ubiquitinated proteins in mammalian cells ^39^ (Fig. 3D). However, expression of IL6, IL8 and MMP3, a set of canonical SASP factors, remained unaffected upon SIRT1 depletion, suggesting that SIRT1 loss itself was insufficient to induce the SASP development (Fig. 3D). We further noticed that DNA damage treatment caused reduction of HSP70, accompanied by increased TSG101, IL8 and MMP3, as evidenced by both transcript and protein assays (Fig. S3A-B). To the contrary, exogenous expression of HSP70 caused decreased levels of TSG101 and ALIX, suggesting a negative correlation between HSP70 expression and sEV production in these cells (Fig. S3C).

**Fig. 3.**
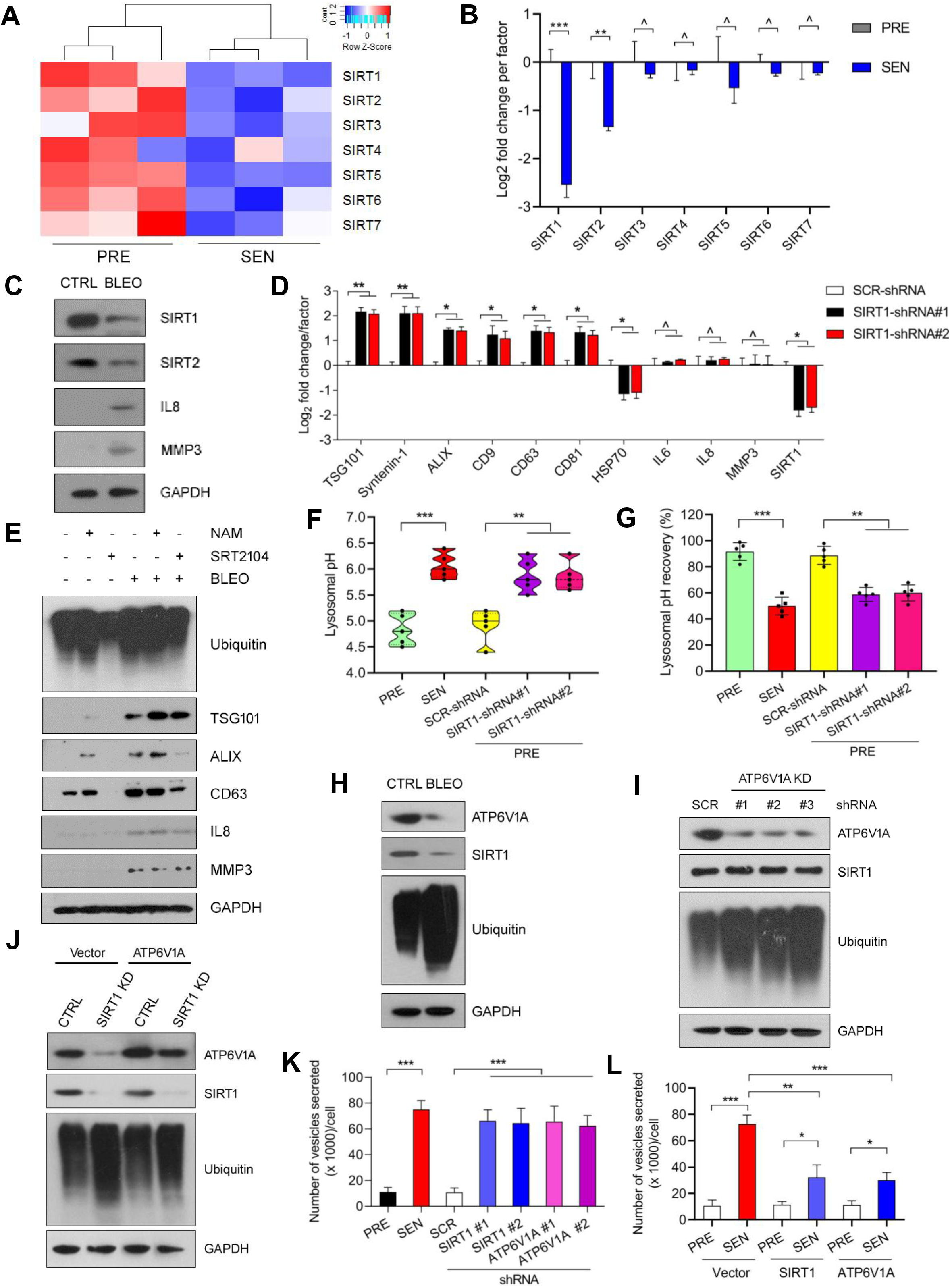
SIRT1 loss mediates lysosomal de-acidification and sEV overproduction through ATP6V1A downregulation in senescent stromal cells. (**A**) Heatmap displaying the expression patterns of human SIRT family members (1-7) in PSC27 cells after senescence. Data derived from RNA-seq. (**B**) Quantitative expression assessment of SIRTs at transcription level in SEN stromal cells by qRT-PCR assays. (**C**) Immunoblot analysis of SIRT1 and the SASP hallmark factors IL8/MMP3 expression in SEN cells induced by bleomycin (BLEO) treatment. (**D**) Transcript analysis of sEV-associated biomarkers upon shRNA-mediated knockdown of SIRT1 in PSC27 cells. (**E**) Stromal cells were subject to treatment by NAM or SRT2104 (a small molecule inhibitor and a small molecule activator, respectively, of SIRT1), alone or together with BLEO, with lysates analyzed by immunoblots after 7 days. (**F**) Lysosomal pH measurements were performed for PRE and SEN cells. Scramble (SCR) and SIRT1-specific shRNAs were transduced to stromal cells to establish stable sublines and assayed for pH, individually. (**G**) Percentage of re-acidification of stromal cell lysosomes posttreatment by Bafilomycin A1. Values were determined with LysoTracker Green DND-26. (**H**) Immunoblot examination of ATP6V1A, SIRT1 and ubiquitin levels in CTRL and BLEO cells. (**I**) Immunoblot analysis of ATP6V1A, SIRT1 and ubiquitin levels in stromal cells transduced with SCR or ATP6V1A-specific shRNAs. (**J**) Immunoblot examination of ATP6V1A, SIRT1 and ubiquitin levels in stromal cells transduced with control or SIRT1-specific shRNAs. In addition, an empty vector or ATP6V1A construct was used for each of these cases before cell lysate collection. (**K**) Number of vesicles released by NTA assays, which were performed on the conditioned media collected from equal numbers of PRE, SEN, and stromal cells transduced with SCR, SIRT1-specific or ATP6V1A-specific shRNAs. (**L**) NTA measurement of vesicle secretion by PRE and SEN stromal cells, in each case an empty vector, SIRT1 or ATP6V1A construct was transduced before collection of conditioned media for NTA assay. ashok, *P* > 0.05; *, *P* < 0.05; **, *P* < 0.01; ***, *P* < 0.001.

The SASP is functionally modulated by several master regulators, including the NF-κB complex, which controls both cell-autonomous and non-cell-autonomous activities of cellular senescence program ^40^. However, whether SIRT1 loss is subject to NF-κB regulation remains unknown. We used Bay 11-7082 (BAY) to treat PSC27 cells, and found that SIRT1 protein level appeared even bounced upon NF-κB suppression, in sharp contrast to many of the SASP factors such as IL8 and MMP3 whose expression was markedly diminished after BAY treatment ^41^ (Fig. S3D). More importantly, NF-κB inhibition substantially reduced expression of TSG101, ALIX, CD63 and CD81 after BLEO treatment, indicating compromised sEV biogenesis even in the setting of DNA damage. However, chemical inhibition of the poly(ADP-ribose) (PAR) polymerase 1 (PARP1), an enzyme functionally involved in cellular senescence, SASP development and NF-κB activation upon DNA damage events ^42^, failed to affect the expression pattern of either SIRT1 or ALIX, suggesting double strand breaks, which underlie ATM- and/or ATR-dependent DNA damage response (DDR), instead of single strand lesions, are indeed responsible for reduced SIRT1 expression and enhanced sEVs biogenesis in DNA-damaged cells (Fig. S3E).

To substantiate the correlation of SIRT1 with sEV production, we used suberoylanilide hydroxamine acid (SAHA) and nicotinamide (NAM), a pan-histone deacetylase (HDAC) inhibitor and a SIRT1-specific inhibitor, respectively, to treat PSC27 cells. Immunoblots indicated that expression of SIRT1, IL8 and MMP3 remained unchanged, HSP70 decreased, while sEV markers including ALIX and CD63 increased when cells were treated by either agent even in the absence of DNA damage, suggesting that SIRT1 activity is critical for the synthesis of sEV-central molecules but not for development of the SASP (Fig. S3F). To the contrary, activation of SIRT1 by SRT2104, a selective SIRT1 activator, substantially diminished sEV signals and protein ubiquitination, alterat ions caused by NAM, regardless of DNA damage treatment, although expression of TSG101, ALIX, CD63, IL8 and MMP3 was remarkably higher when cells were exposed to BLEO (Fig. 3E). Thus, SIRT1 is a critical but negative modulator of sEV biogenesis, and functionally responsible for restrained proteome-wide ubiquitination.

Enhanced production or reduced degradation of ubiquitinated proteins by the proteasome or autophagy can result in elevated amount of ubiquitinated proteins ^39^. To assess proteasome integrity in damaged cells, we used the proteasome inhibitor MG132 to treat PSC27, and found pronouncedly increased protein ubiquitination after genotoxic exposure (Fig. S3G). Upon treatment with Bafilomycin A1, a selective inhibitor of the late phase of autophagy that prevents maturation of autophagic vacuoles by targeting vacuolar H+-ATPases (V-ATPases) and restraining fusion between autophagosomes and lysosomes ^43^, we noticed elevated levels of autophagy markers p62 and LC3 II, regardless of cell exposure to genotoxic stress, indicating the integrity of autophagy even with SIRT1 loss and protein ubiquitination in these senescent cells (Fig. S3H). Upon treatment by cycloheximide (CHX), a protein synthesis inhibitor, we observed markedly reduced protein ubiquitination, but not SIRT1 level (Fig. S3I). The total protein of EGFR, a transmembrane receptor subject to polyubiquitination and proteasome-mediated degradation ^44^, remained largely unaffected when BLEO-damaged cells were exposed to CHX. Indeed, the relative constant amount of EGFR protein was in line with reduced level of ubiquitination in CHX-treated cells, which presumably had decreased capacity in mediating protein turnover. Thus, SIRT1 was not subject to significant protein degradation in senescent cells, with its decrease mainly attributable to downregulation of SIRT1 starting from the transcription level, although accumulation of ubiquitinated proteins implies functional deficiency of lysosomes.

We next interrogated whether SIRT1 loss affects lysosomal function by assaying the pH of lysosomes with LysoSensor yellow/blue dextran ratiometric probe, which accumulates in acidic organelles as the result of protonation. In contrast to PRE cells whose lysosome pH was 4.8, consistent with the reported pH of physiologically normal lysosomes ^45^, SEN cells exhibited a lysosomal pH approaching 6.0 (Fig. 3F). To determine whether the increase of SEN cell lysosomal pH results from a defective proton pump, we performed lysosomal re-acidification assays by first enhancing the pH by Bafilomycin A1-treatment, then adding LysoTracker to assess the rate of lysosomal pH recovery. While the pH of PRE cell lysosomes basically recovered within 60 min, the pH recovery of SEN cell lysosomes appeared markedly slower (Fig. 3G). Interestingly, upon SIRT1 depletion from these cells, a similar tendency of lysosomal pH increase and post-stress pH recovery deficiency was observed, which largely resembled that of SEN cells (Fig. 3F-G).

As the data suggested defective proton pump in SEN cells, we analyzed V-ATPases, a set of multi-subunit enzymes functionally supporting the acidification of late endosomes and lysosomes^46^. Immunoblots showed that expression of subunit A of V1 (ATP6V1A) substantially decreased upon cellular senescence, in parallel with SIRT1 reduction and pan-ubiquitination (Fig. 3H). Further analysis showed enhanced ubiquitination when ATP6V1A was eliminated, even with SIRT1 level unaffected, suggesting ATP6V1A is responsible for controlling protein ubiquitination, a process downstream of SIRT1 regulation (Fig. 3I). While SIRT1 elimination remarkably enhanced ubiquitination, ectopic expression of ATP6V1A was able to rescue such an effect (Fig. 3J). We then assessed the capacity of sEV secretion, and found ATP6V1A knockdown caused a significant increase in the number of sEVs released from stromal cells, an effect that largely resembled that of SIRT1 depletion (Fig. 3K). To the contrary, overexpression of either SIRT1 or ATP6V1A prominently restrained sEV production in SEN cells, although no changes were observed in PRE cells (Fig. 3I). Thus, our data consistently demonstrated that SIRT1 loss causes lysosomal de-acidification and is responsible for sEV overproduction via ATP6V1A downregulation in SEN cells, a mechanism reported for certain cancer cell types ^35^.

### Senescent stromal sEVs enhance the malignancy of recipient cancer cells particularly therapeutic resistance

We next asked whether sEVs secreted by SEN stromal cells can exert profound impacts on cancer cells. We generated a stable PSC27 subline (PSC27-CD63) with a construct (pCT-CD63-eGFP), permitting to utilize the eGFP-fusion CD63 to mark cellular compartment, organelles and structures to enable long term and in-depth tracing of sEV biosynthesis, secretion, and uptake. Upon collection of PSC27-derived sEVs, we applied them directly to treat prostate cancer (PCa) cells in culture (Fig. 4A). Not surprisingly, eGFP-marked CD63 was easily detected, although mainly in the perinuclear region of PCa cells, suggesting active uptake of these vesicles by recipient cancer cells (Fig. 4B).

**Fig. 4.**
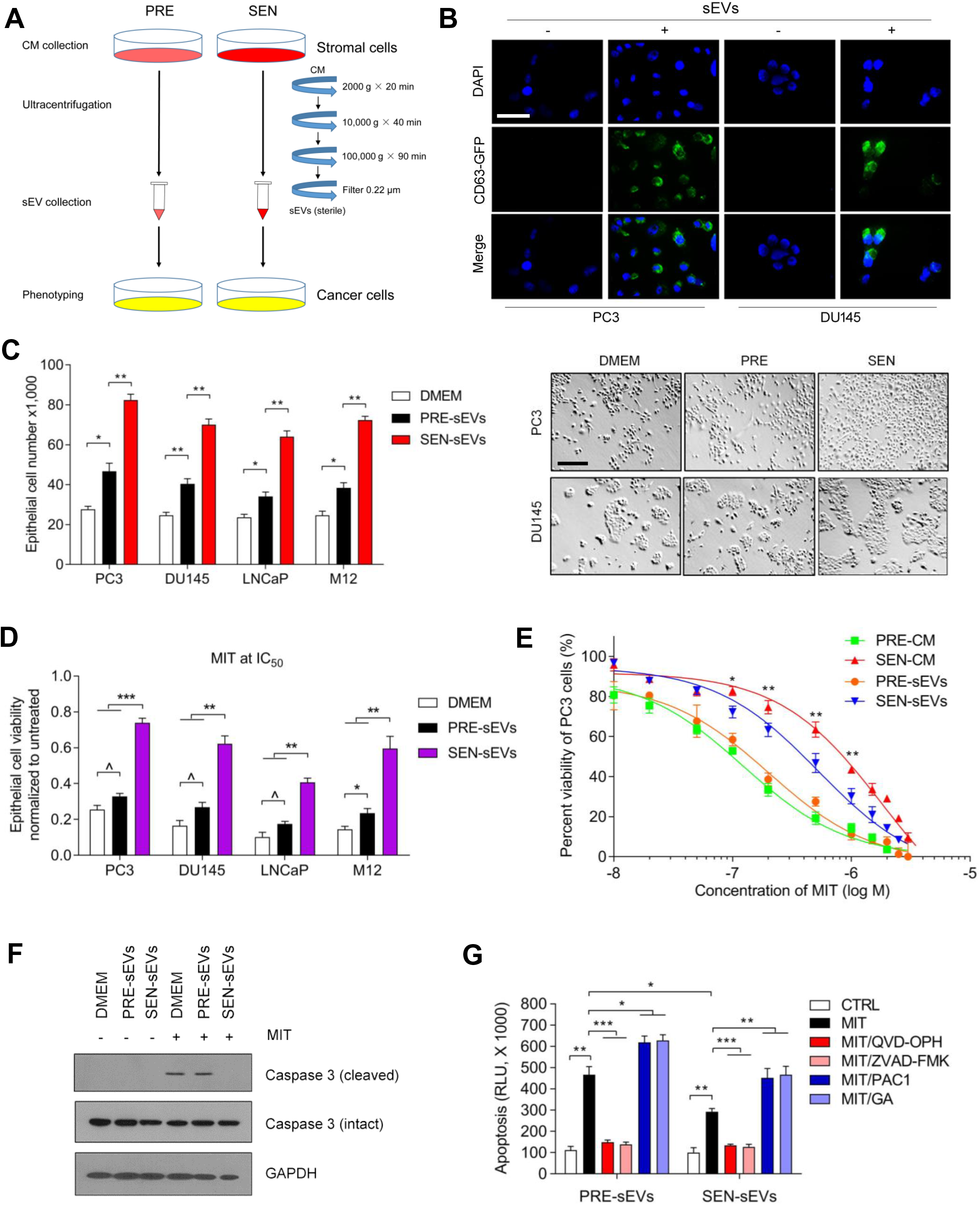
Senescent stromal sEVs promote the malignant phenotypes particularly drug resistance of prostate cancer cells. (**A**) Schematic workflow of conditioned media (CM) collection, sequential ultracentrifugation-based sEV preparation from stromal cells and *in vitro* phenotyping of cancer cells. (**B**) PSC27 cells were transduced with the pCT-CD63-eGFP construct, and the secreted sEVs were collected to treat PC3 and DU145 cells prior to immunofluorescence imaging. Scale bar, 20 μm. (**C**) PCa cells were treated with sEVs from PSC27 cells for 3 d, and subject to proliferation assay. DMEM, blank control; PRE-sEVs, sEVs of PRE PSC27 cells; SEN-sEVs, sEVs of SEN PSC27 cells. Right, representative images. Scale bar, 40 μm. (**D**) Chemoresistance assay of PCa cells cultured with stromal cell sEVs described in (C). Mitoxantrone (MIT) was applied at the concentration of IC50 value pre-determined per cell line. Cells were cultured in DMEM, which was also used as a drug vehicle. (**E**) Dose-response curves (non-linear regression/curve fit) plotted from drug-based survival assays of PC3 cells cultured with the sEVs collected from PRE or SEN PSC27 cells, and concurrently exposed to a wide range of MIT concentrations. (**F**) Immunoblot analysis of caspase 3 cleavage in PC3 cells upon exposure to different conditions. MIT was applied at the IC50 value predetermined for PC3. (**G**) Apoptotic assay for combined activities of caspase 3/7 determined 24 h after exposure of PC3 cells to stromal cell sEVs. Cancer cells were treated by MIT in the presence or absence of caspase inhibitors including QVD-OPH and ZVAD-FMK, or caspase activators including PAC1 and gambogic acid (GA). *, *P* < 0.05; **, *P* < 0.01; ***, *P* < 0.001.

Subsequent *in vitro* assays demonstrated that SEN stromal sEVs can significantly enhance the proliferation of several PCa cell lines including PC3, DU145, LNCaP and M12, in contrast to sEVs released from PRE stromal cells, which though also generated detectable effects (Fig. 4C). We further observed markedly increased migration and invasion of these cells upon exposure to SEN stromal sEVs (Fig. S4A-B). More importantly, PCa cells showed significantly enhanced resistance to mitoxantrone (MIT), a type II topoisomerase inhibitor administered to treat several types of malignancies including PCa in clinical medicine (Fig. 4D). Although PRE stromal sEVs did not confer significant benefits on the survival of the majority of PCa cell lines we assayed, SEN stromal sEVs pronouncedly enhanced the resistance of cancer cells to MIT-induced cytotoxicity (Fig. 4D). We further assessed the influence of stromal sEVs on cancer cell survival with MIT, which was specifically prepared in a range of doses (0.01 ∼ >1.0 μM) designed to resemble its serum concentrations in clinical conditions. The data suggested that PRE stromal sEVs did not significantly change PC3 survival when cells were exposed to MIT, whereas SEN stromal sEVs did (Fig. 4E). We noticed that the two groups of stromal sEVs differed most dramatically in the range of 0.1-1.0 μM MIT, although further augment of drug concentration consistently resulted in rapid clearance of cancer cells presumably due to cell intolerance against overwhelming cytotoxicity.

Mechanistic dissection revealed that MIT induced cleavage of caspase 3 in cancer cells, a process that was modestly affected by PRE stromal sEVs but remarkably weakened by SEN stromal sEVs (Fig. 4F). Although alternative cell survival pathways may be operative, our data suggest that SEN stromal sEVs drive cancer resistance likely via a caspase-counteracting mechanism. We further applied QVD-OPH and ZVAD-FMK, two potent pan-caspase inhibitors, as well as PAC1 and gambogic acid (GA), two typical caspase activators, to individually treat PC3 cells shortly before MIT exposure. Cell apoptosis was substantially attenuated in the presence of QVD-OPH or ZVAD-FMK (*P* < 0.001) (Fig. 4G). However, once the procaspase-activating compound PAC1 or GA was used, apoptosis index was markedly elevated, thus offsetting the anti-apoptosis effect of SEN stromal sEVs (*P* < 0.01). The data were largely reproduced when docetaxel (DOC), another chemotherapeutic drug that interferes with microtubule depolymerization, was applied to the system (Fig. S4C-D). Thus, our results consistently demonstrate that SEN stromal sEVs restrains caspase-dependent apoptosis of cancer cells, a process that underlies its resistance-boosting capacity via nanoparticle secretion-based paracrine influence on cells exposed to cytotoxic agents.

### Expression profiling of prostate cancer cells is subject to profound alteration by senescent stromal sEVs

Given the remarkable changes of PCa cell phenotypes induced by senescent stromal sEVs, we next sought to determine their influence on the expression pattern of cancer cells. We first chose to perform RNA-seq to quantitate gene expression modifications and profile the transcriptomics after treatment of PCa cells with SEN stromal sEVs. Bioinformatics output showed that 344 transcripts were upregulated or downregulated significantly (≥ 2-fold, *P* < 0.05) in PC3 cells (Fig. S5A-B), while the expression of 422 transcripts was modified in SEN stromal sEV-affected DU145 cells (Fig. S5C-D). Among the upregulated gene products, many are correlated with tumor development, such as CEACAM5, KRT15, IFI6, KIF20A, CEMIP and SERPINB3 in PC3, or DHRS2, IL21R, FGFBP1, DKK1, PRSS2 and KLK6 in DU145 (Fig. S5E-F). We noticed that these proteins are mostly involved in key biological processes such as signal transduction, cell communication, cell metabolism, energy pathways, cell growth and maintenance (Fig. S5G-H). Therefore, data from bioinformatics analysis generally support the findings derived from *in vitro* assays, which showed enhanced cell proliferation, migration and invasiveness activities of cancer cells upon exposure to SEN stromal sEVs (Fig. 4C and Fig. S4A-B).

Next, we performed comparative assessment by scrutinizing the genes whose expressed was co-upregulated in both PC3 and DU145 cells (≥ 2-fold, *P* < 0.05). The data presented 22 genes that fell in this category, with 199 and 169 genes being uniquely upregulated in PC3 and DU145, respectively (Fig. S5I). Interestingly, we noticed the ATP binding cassette subfamily B member 4 (ABCB4), a full transporter and member of the p-glycoprotein family of membrane proteins with phosphatidylcholine as its substrate, showed up at the top of the PC3-DU145 co-upregulated gene list (Fig. S5J). GO analysis indicated that ABCB4 is involved in multiple activities including glycoside transport, carbohydrate export, response to drug, fenofibrate, external biotic stimulus and cellular hyperosmotic salinity (Fig. S5K). Alternatively, KEGG appraisal underscored its biological implications as a typical ABC transporter (Fig. S5L).

After mapping ABCB4-involved protein-protein interaction (PPI) curated with the STRING database interactome ^47^, we generated a network highlighting the interaction of ABCB4 with a handful of proteins such as PPARA, FABP1, CREBBP, CHD9 and CARM1 in human cells (Fig. S5M). The PPI network encompassed protein target nodes (*n* = 11) connected by edges (*n* = 55) with an average node degree of 10 (local clustering coefficient of 1, PPI enrichment *P* < 1.0e-16). We calculated the evidence based on GO and other databases including KEGG, PFAM and InterPro, with binary human PPI data and enrichment results validated (Table S4).

### Senescent stromal sEVs promote therapeutic resistance by upregulating ABCB4 expression in recipient cancer cells

Given the substantial impact of SEN stromal sEVs on transcriptome-wide expression of recipient cancer cells, we sought to explore the mechanism(s) supporting stromal sEV-enhanced malignancies particularly drug resistance of PCa cells. RNA-seq data showed that upon treatment with SEN stromal sEVs, PC3 and DU145 cells exhibited significantly (*P* < 0.05 for both lines) upregulated expression of ABCB4, while other members of the ABCB subfamily remained largely unchanged (Fig. 5A-B). Subsequent transcript assays essentially supported these results (Fig. 5C-D). We further confirmed the SEN stromal sEV-inducible pattern of ABCB4 by immunoblots (Fig. 5E).

**Fig. 5.**
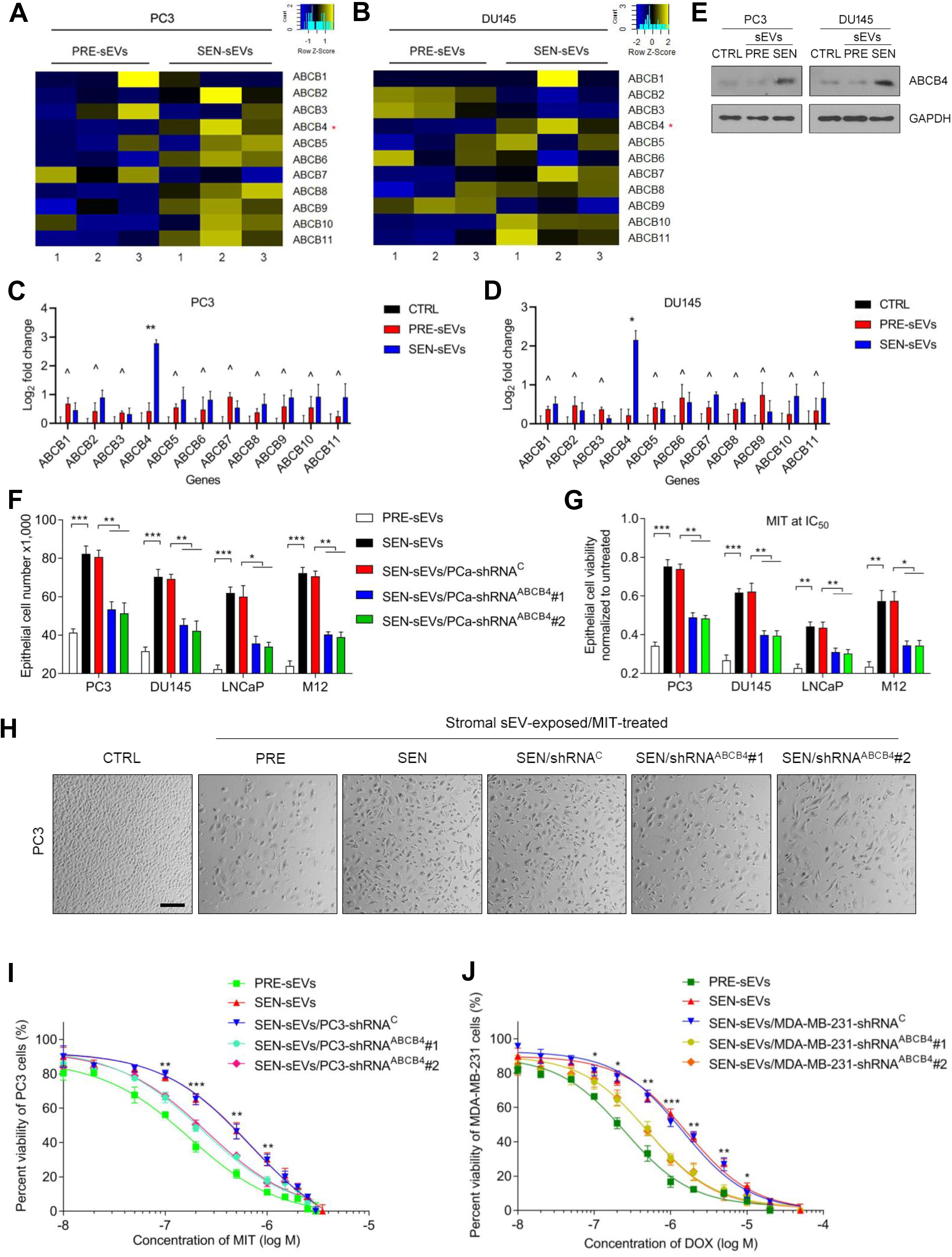
Senescent stromal sEVs enhance drug resistance via upregulation of ABCB4 in recipient cancer cells. (**A**) Heatmap depicting the expression patterns of 11 members of human ABCB subfamily in PC3 cells upon exposure to SEN vs PRE stromal sEVs. (**B**) Heatmap displaying the expression profiles of 11 members of the ABCB subfamily in DU145 cells upon exposure to SEN vs PRE stromal sEVs. (**C**) Quantitative RT-PCR transcript assays of the expression of human ABCB subfamily members in PC3 cell line after incubation with sEVs for 3 consecutive days. Signals were normalized to the control group (DMEM). (**D**) Transcript expression assays of human ABCB subfamily member expression in DU145 cell line. (**E**) Immunoblot analysis of ABCB4 expression in PC3 and DU145 cells after exposure to SEN vs PRE stromal sEVs for 3 consecutive days. (**F**) Proliferation assay of PCa cells exposed to SEN vs PRE stromal sEVs for 3 days, with cancer cells depleted of ABCB4 via shRNA-mediated knockdown. (**G**) Chemoresistance assay of PCa cells exposed to sEVs from SEN vs PRE stromal cells. ABCB4 was eliminated in cancer cells, with MIT applied at the concentration of IC50 value pre-determined per cell line. (**H**) Representative images of PC3 cells examined under conditions described in G. Scale bar, 100 μm. (**I**) Dose-response curves (non-linear regression/curve fit) plotted from drug-based survival assays of PC3 cells exposed to stromal sEVs, and concurrently treated by a wide range of MIT concentrations. (**J**) Dose-response curves of MDA-MB-231 cells similar to those shown in I, with doxorubicin (DOX)-based survival assays performed. ashok, *P* > 0.05; *, *P* < 0.05; **, *P* < 0.01; ***, *P* < 0.001.

Next, to address the relevance of ABCB4 upregulation to cancer cells, we eliminated this multidrug resistance protein of an ATPase-associated function responsible for drug efflux with gene-specific shRNAs. Of note, the gain of function in cell proliferative rate after exposure to SEN stromal sEVs was largely diminished upon depletion of ABCB4 from representative PCa cell lines (Fig. 5F). We observed significantly weakened migration and invasion when ABCB4 was eliminated from these cells (Fig. S6B-C). The effects caused by ABCB4 removal from cancer cells were further observed in chemoresistance assays, which showed a similar tendency among the PCa cell lines examined at their individual MIT concentration of IC50 (Fig. 5G-H). To expand the findings, we further examined cell survival capacity across a wide window of MIT concentrations with PC3, and found ABCB4 knockdown caused a remarkable reduction in cell survival upon treatment by MIT in the range of 0.1 ∼ 1.0 μM (Fig. 5I). Thus, our data consistently support the key function of ABCB4 in mediating resistance to a chemotherapeutic agent, a property acquired upon uptake of SEN stromal sEVs by cancer cells.

To validate the generality of these findings, we collected sEVs from HBF1203, the breast stromal cell line, which enters cellular senescence upon DOX treatment (Fig. S1E-H). Following a similar procedure, we eliminated ABCB4 from MDA-MB-231, a breast cancer cell line, before exposure to SEN stromal sEVs. The data from breast stroma-cancer assessments closely resembled those derived from prostate stroma-cancer assays, by showing the effect of ABCB4 depletion on BCa cell survival in a DOX concentration range of 0.1 ∼ 10 μM which approaches its plasma level of BCa patients in clinics ^48^ (Fig. 5J).

### Activating SIRT1 restrains sEV production by senescent cells and promotes anticancer efficacy

Since SIRT1 expression is reduced in SEN stromal cells, a process responsible for increased sEV production, we reasoned the possibility of minimizing sEV production by SEN stromal cells though targeting SIRT1. To this end, we chose SRT2104 and SRT1720, two potent selective activators of SIRT1, to treat stromal cells. Simultaneous exposure of PSC27 cells to BLEO and either SIRT1 activator significantly decreased the number of sEVs released by SEN PSC27 cells (Fig. 6A and Fig. S7A). More importantly, we observed markedly declined ABCB4 expression and diminished chemotherapeutic resistance of PCa cells (PC3 as a representative) upon incubation with sEVs secreted by SEN stromal cells which were simultaneously exposed to BLEO-delivered genotoxicity and a SIRT1 activator (SEN/SRT2104 or SEN/SRT1720), in contrast to the senescence-naive group (SEN or SEN/DMSO) (Fig. S7B and Fig. 6B). The data derived from PSC27 and PC3 were largely reproducible by a similar set of assays performed with HBF1203 and MDA-MB-231, both lines of human breast origin (Fig. S7C-D). Thus, targeting SIRT1 with pharmacological activators can restrain sEV production, a treatment strategy that is effective in governing the phenotype of SEN stromal cells but indeed correlated with significantly weakened drug resistance of cancer cells.

**Fig. 6.**
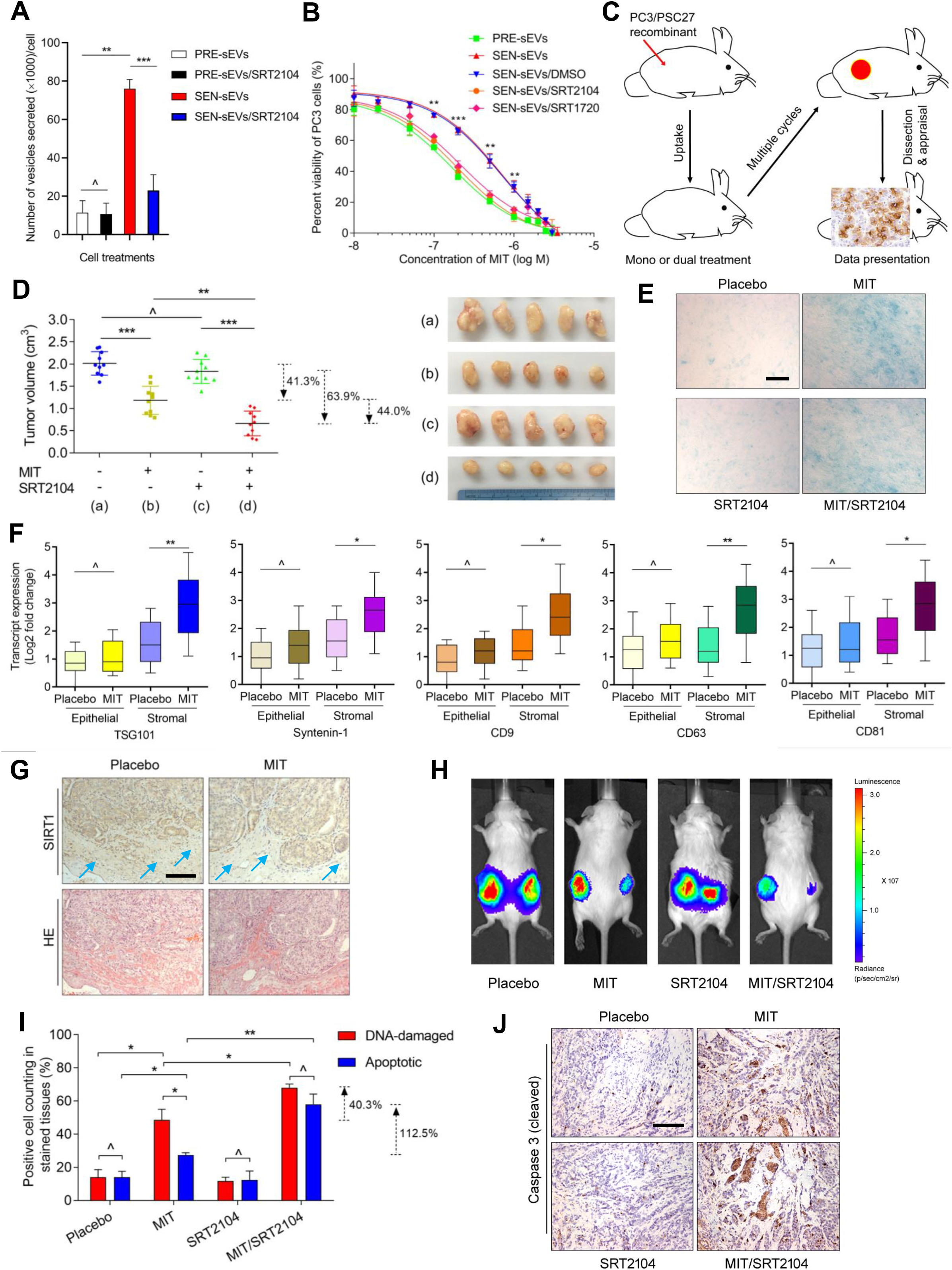
Targeting SIRT1 with an agonist compound to control sEV production by senescent stromal cells improves chemotherapeutic efficacy. (**A**) Activation of SIRT1 restrains sEV production in SEN stromal cells. SRT2104, a selective SIRT agonist, was used to treat PSC27 cells alone or together with BLEO for 7-10 d. (**B**) Dose-response curves plotted from drug-based survival assays of PC3 cells exposed to sEVs derived from stromal cells treated by BLEO and/or SRT2104 (or SRT1720, another selective SIRT1 agonist). (**C**) Schematic illustration of preclinical treatments performed with immunodeficient mice. (**D**) Statistics of tumor end volumes. PC3 cells were xenografted together with PSC27 cells (totally 1.25 × 10^4^, in a cancer/stromal ratio of 4:1) to the hind flank of mice. SRT2104 and the chemotherapeutic agent MIT was administered alone or together to induce tumor regression. (**E**) Representative images of *in vivo* cellular senescence after SRT2104- and/or MIT-mediated treatment. Scale bar, 100 μm. (**F**) Transcript assay of sEV biomarkers expressed in epithelial and stromal cells isolated from the tumors of experimental mice. (**G**) Representative IHC images of SIRT1 expression in tissues isolated from placebo or MIT-treated animals. Scale bar, 200 μm. (**H**) Representative bioluminescence images (BLI) of PC3/PSC27 tumor-bearing animals in the preclinical trial. (**I**) Statistical evaluation of DNA-damaged and apoptotic cells in the biospecimens. Values are presented as percentage of cells positively stained by IHC with antibodies against γ-H2AX or caspase 3 (cleaved). (**J**) Representative IHC images of caspase 3 (cleaved) in tumors at the end of therapeutic regimes. Scale bar, 200 μm. ashok, *P* > 0.05; *, *P* < 0.05; **, *P* < 0.01; ***, *P* < 0.001.

Given the prominent efficacy of SIRT1 activator-mediated control of sEV production by SEN stromal cells, we asked whether the *in vitro* findings can be technically repeated by *in vivo* studies. To precisely address tumor-stroma interactions in a TME context with structural and functional integrity, we generated tissue recombinants by admixing PSC27 with PC3 at a pre-optimized ratio before subcutaneously injecting them to the hind flank of experimental mice with severe combined immunodeficiency (SCID). Chemotherapeutic agent MIT and SRT2104 (chosen for preclinical studies due to its slightly higher efficacy than SRT1720; SRT2104 currently in phase 2 trials, while SRT1720 failed) were administered as mono or dual agents starting from the 3^rd^ week post tumor implantation, after which animals received treatment in a metronomic manner to mimic clinical conditions (Fig. 6C and Fig. S7E). Data from endpoint measurement of tumor size indicated that MIT administration caused remarkably delayed tumor growth, validating the efficacy of MIT as a cytotoxic agent (41.3%, *P* < 0.001) (Fig. 6D). Although treatment with SRT2104 did not significantly change tumor volume, co-administration of MIT and SRT2104 caused maximal reduction of tumor mass (63.9% shrinkage in contrast to SRT2104, or 44.0% decrease in relative to MIT given in the mono-treatment) (Fig. 6D). We observed substantially enhanced cellular senescence in tumors dissected from animals that underwent therapeutic regimens involving MIT, regardless of SRT2104 administration (Fig. 6E).

Treatment-induced cellular senescence was accompanied by in-tissue development of the SASP, as evidenced by increased expression of SASP hallmark factors such as IL6, IL8, IL1α, MMP3 (Fig. S7F). Although stromal cells exhibited consistently increased SASP expression, cancer epithelial cells did not show a typical senescence tendency, including expression of p16 which remained largely unchanged after treatment (Fig. S7F). This was likely due to the emergence of treatment-resistant cancer cells and expansion of resistance colonies during the chemotherapeutic regimen, a process responsible for tumor relapse posttreatment. Importantly, we observed significantly decreased signals of SIRT1 in stromal cells post-treatment, which was in line with our *in vitro* data (Fig. 3A-C). Reduction of SIRT1 in stroma was accompanied by consistently enhanced expression of sEV markers including TSG101, Syntenin-1, CD9, CD63 and CD81 (Fig. 6F). In contrast, cancer cells did not seem to develop such a distinct pattern.

Loss of SIRT1 expression was also revealed by immunohistochemistry (IHC) staining of mouse tissues. In contrast to stromal cells which showed a generally declining tendency, adjacent cancer cells appeared largely unchanged in SIRT1 expression (Fig. 6G). The data suggested the presence of a differential expression mechanism that demarcates stromal cells from their cancer epithelial counterparts. IF staining of tissues indicated that there was a remarkable increase of ABCB4 in cancer cells, but not stromal cells, after exposure of animals to MIT, although the signals were minimized when SRT2104 was delivered alongside MIT (Fig. S7G-H).

As sEVs can foster pre-metastatic niche formation and mediate distant metastasis of cancer cells ^49, 50^, we interrogated whether there are metastatic events in these animals. Bioluminescence imaging (BLI) of xenografts generated with PC3 cells stably expressing luciferase (PC3-luc) and PSC27 stromal cells excluded activities of cancer cell dissemination from the primary sites, while the relative BLI signal intensities essentially supported tumor growth patterns observed in xenografted mice (Fig. 6H). The data suggest that classic chemotherapy combined with a SIRT1-activating agent can achieve tumor regression more effectively than chemotherapy alone, a therapeutic strategy that holds a prominent potential to circumvent the SEN stroma-induced pathological exacerbation, specifically acquired resistance.

Subsequent tissue assessment indicated that MIT administration caused dramatically increased DNA damage and apoptosis, as evidenced by DDR foci measurement and caspase 3 cleavage appraisal (Fig. 6I). Although SRT2104 alone did not induce typical DDR or apoptosis, significantly elevated indices of both activities were observed in tissues of animals that underwent MIT/SRT2104 combinatorial treatment. In contrast to the MIT-only group, there was a pronounced increase of DDR and apoptosis in MIT/SRT2104 animals (40.3% and 112.5%, respectively), suggesting a remarkable potential of SRT2104 in promoting cancer cell clearance and achieving tumor regression when synergized with classic chemotherapy *in vivo* (Fig. 6I-J).

### SIRT1 decline in senescent stromal cells and ABCB4 expression in cancer cells predict adverse outcome post-chemotherapy

We next interrogated the pathological relevance of SIRT1 reduction in stroma and ABCB4 elevation in cancer cells to patient survival after chemotherapy. Upon IHC staining, we observed enhanced p16^INK4a^ expression in stromal cells of patients who experienced chemotherapeutic intervention, while most cancer cells remained largely negative (Fig. 7A). The data revealed comprehensive occurrence of cellular senescence in the stroma while cancer cells seemingly resisted in the gland, presumably through expansion from colonies that survived anticancer treatments via development of drug resistance. We noticed remarkably decreased expression of SIRT1 in stromal, rather than their adjacent cancer cells, although the latter showed elevated ABCB4 expression in posttreatment patients (Fig. 7A).

**Fig. 7.**
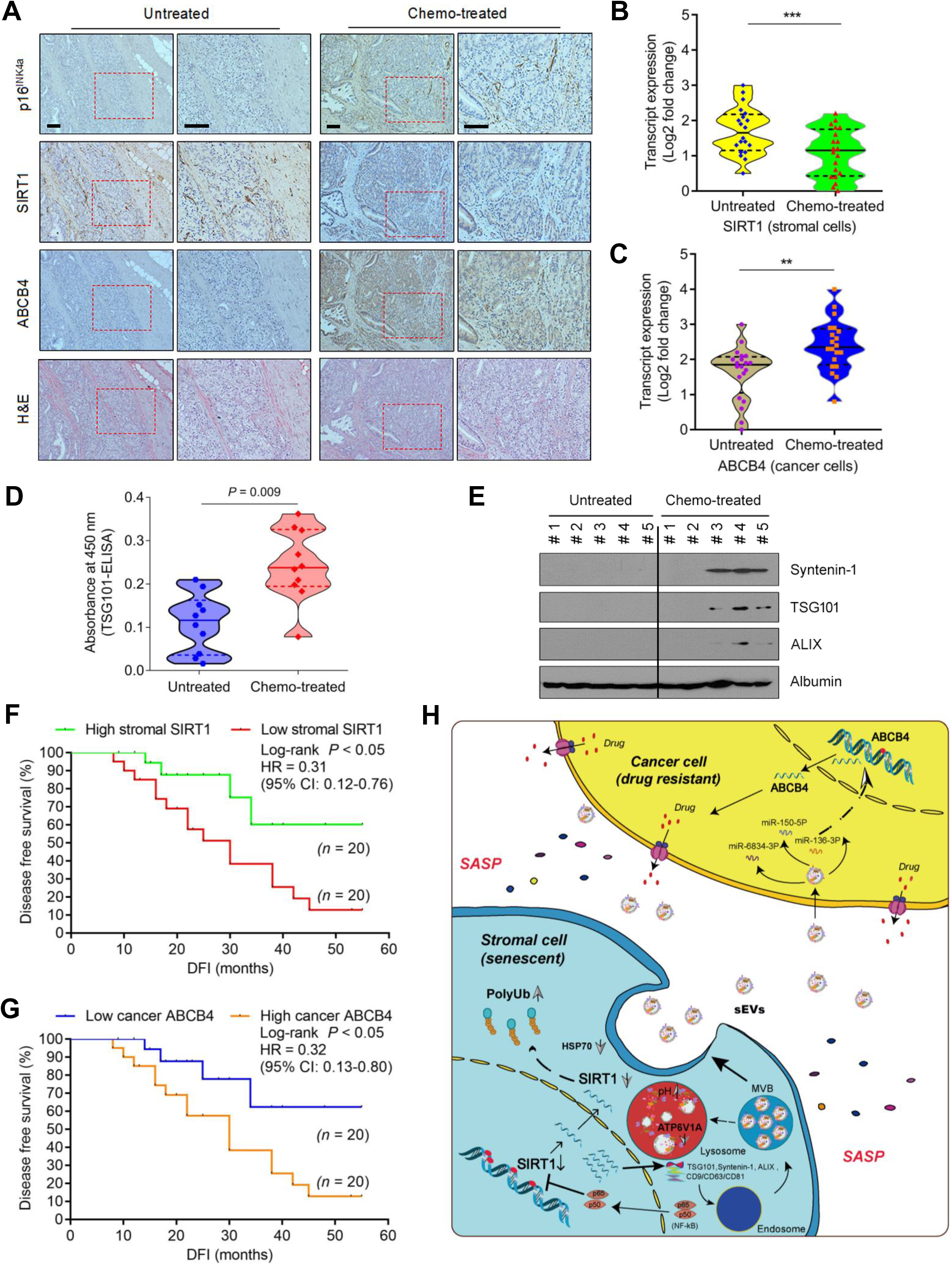
SIRT1 loss in stroma and ABCB4 upregulation in cancer cells predict adverse clinical outcome. (**A**) IHC assessment of human prostate tumors before and after chemotherapy. From top to bottom, representative images of p16^INK4a^, SIRT1 and ABCB4 expression in biospecimens of human PCa patients. All samples from a longitudinal study of a same patient. Scale bars, 200 μm. (**B**) Transcript expression assay of SIRT1 in stromal cells isolated via LCM from tumor samples (20 untreated and 20 chemo-treated PCa patients randomly selected). (**C**) Transcript expression assay of ABCB4 in cancer cells isolated via LCM from tumor samples. (**D**) Measurement of sEVs in peripheral blood of PCa patients with TSG101-specific ELISA performed among 10 patients randomly selected from each of untreated and chemo-treated cohorts. (**E**) Immunoblot of Syntenin-1, TSG101, and ALIX, a set of typical markers of sEVs, in serum of PCa patients collected before and after chemotherapy. Albumin, loading control for patient serum samples. (**F**) Kaplan-Meier (KM) survival analysis of chemo-treated PCa patients stratified with SIRT1 levels in tumor stroma. (**G**) KM assay of chemo-treated PCa patients stratified with ABCB4 levels in tumor foci. (**H**) Illustrative working model of the impact of senescent stromal sEVs on development of therapeutic resistance of cancer cells in a TME niche.

Upon precise isolation of stromal and cancer cell subpopulations with laser capture microdissection (LCM) and evaluation by qRT-PCR, we noticed SIRT1 reduction and ABCB4 elevation in stromal and epithelial cells, respectively (Fig. 7B-C). Thus, transcript expression data were essentially consistent with those acquired from tissue staining appraisal of these clinical specimens.

We further performed a sample-based clinical investigation by randomly choosing two subgroups of PCa patients (10 for each) from the overall cohort, including one that did not undergo chemotherapy and the other that experienced chemotherapy, with patient-derived peripheral blood samples available from both subgroups. TSG101-specific ELISA tests suggested significantly enhanced levels of circulating sEVs in the serum of post-treatment patients, as compared with those untreated individuals (Fig. 7D). Surprisingly, immunoblots even showed clearly visible signals of circulating sEVs (Syntenin-1, TSG101 and ALIX-positivity) in the serum of post-treatment patients (3 out of 5 cases), in sharp contrast to samples from those untreated which were generally negative (Fig. 7E). Therefore, sEVs in the circulating system of post-treatment patients hold the potential to be evaluated as blood-borne substance for clinical monitoring via examination with routine biotechniques.

We next sought to establish the consequence of enhanced sEV biogenesis in these patients. After tissue staining against SIRT1 and ABCB4, we assessed these patients by associating the results with their survival. Strikingly, there was a significant and positive correlation between stromal SIRT1 expression and patient disease-free survival (DFS) in the post-treatment PCa cohort (Fig. 7F). To the contrary, a significant but negative correlation was observed between cancer cell ABCB4 expression and patient survival (Fig. 7G). Thus, SIRT1 decline-mediated sEV production and release by stromal cells, combined with ABCB4 upregulation in cancer cells, can provide a novel and precise avenue for prognosis of advanced malignancies in future cancer medicine. Specifically, given the prominent relevance of SIRT1 loss in sEV biogenesis of SEN stromal cells and its hitherto well documented pathological implications in human pathologies, we would propose SIRT1 as a functionally multifactorial target in clinical oncology.

## DISCUSSION

Mutual interaction between cancer cells and the surrounding TME is crucial for malignant progression. Numerous studies have unraveled the paracrine signaling of various cytokines, chemokines and growth factors through their receptors as a key means of intercellular communication in the microenvironmental niche ^51, 52^. In contrast, EVs have recently emerged as another mechanism to mediate cell-cell interactions, which can be secreted from various cell types to generate a unique TME context ^53^. The sEVs are a subclass of EVs and attract increasing attention as they contain various proteins, mRNAs, microRNAs, lipids and even DNA fragments to target cells, thereby altering their phenotypes. Technically, sEVs can be distinguished from the other major type of EVs, namely the MVs, based on their average size and biogenesis. MVs range from 0.2 to 2.0 μm in diameter and directly bud off from plasma membrane, whereas sEVs (in some literatures, alternatively designated as exosomes) are usually ∼30-150 nm (< 200 nm) in size and packed within MVBs, which release sEVs into the extracellular space upon fusion with cell membrane ^54, 55^. Despite the exponential growth of the EV field, a great deal remains unclear about the mechanisms regulating EV production, and most cancer-associated EV studies so far focus on EVs derived from cancer cells or normally proliferating cells. In contrast, here we present evidence to show that senescent stromal cells (including PSC27 and HBF1203, each of human origin), which abundantly reside in host tissues and closely reflect microenvironmental conditions in the TME, are responsible for cancer progression by producing numerous sEVs to shape drug resistance acquired from the treatment-damaged TME (Fig. 7H).

Sirtuins (SIRT1-7) belong to the third class of HDACs, whose activities are dependent on NAD^+^ ^56^. In this study, we revealed SIRT1 as the most dramatically declined sirtuin molecule in senescent stromal cells, with accumulation of ubiquitinated proteins in cytosol. Protein quality control is critical for maintenance of cell physiology and tissue homeostatis, while damaged proteins need to be restored or degraded, a process mainly supported by molecular chaperones and the ubiquitin-proteasome system. Interestingly, both proteasome and autophagy machineries remained largely intact in these cells, factors experimentally excluded from affecting SIRT1 protein level. Although reasoned to be subject to regulation at transcription level, SIRT1 reduction eventually causes enhanced expression of multiple sEV markers including TSG101, Syntenin-1, ALIX and tetraspanins, functionally promoting sEV biogenesis in senescent stromal cells. Simultaneously, SIRT1 loss is responsible for ATP6V1A downregulation, resulting in defective lysosomal acidification and sEV overproduction by senescent cells. The data are reminiscent of a recent study which demonstrated that genetic or pharmacological inhibition of SIRT1 destabilizes the mRNA of the lysosomal V-ATPase proton pump and causes its downregulation (ATP6V1A), leading to deficient lysosomal acidification, MVB enlargement and enhanced exosomes production ^35^. While our observation was mainly regarding senescent stromal cells, distinct from the discovery recently made in human breast cancer cells, a similarly occurring MVB expansion revealed by our *in vitro* assays suggests that senescent stromal cells somehow resemble cancer cells with regard to sEV biogenesis. Further, SIRT1 loss-driven ATP6V1A downregulation results in defective lysosomal acidification and a manifest increase of pH to approximately 6.0, partially explaining the comprehensively observed SA-β-Gal staining positivity of senescent cells, a phenomenon reported long time ago but rarely elucidated well for the underlying mechanism(s) ^57^.

There is a growing interest in the discovery of small molecules modifying sirtuin activities, although most sirtuin activators have been described intensively for SIRT1 ^58^. Specifically, SIRT1 has been suggested to play multiple and, in some cases, contradictory roles in cancer, as that SIRT1 can function as either a tumor suppressor or tumor promoter, depending on the tissue type and pathological context ^35^. *In vivo* stimulation of SIRT1 either with agonist or NAD^+^ precursors can extend the lifespan of mice and protect against diverse aging-related diseases ^59^. Resveratrol is a natural polyphenol compound which activates SIRT1, holding potential in treatment or prevention of tumorigenesis and aging-related diseases, but it has limited bioavailability ^60^. Molecules structurally unrelated to resveratrol, including SRT1720 and SRT2104, are typically sirtuin-activating compounds (STACs) developed to stimulate sirtuin activities more potently than resveratrol, with markedly improved bioavailability ^61^. In our study, we disclosed that SRIT1 activators such as SRT2104 can effectively restrain sEV biogenesis in senescent stromal cells. Although cancer cells exposed to the stromal CM containing sEVs developed remarkably enhanced malignancy particularly drug resistance, a process principally mediated by ABCB4 as a p-glycoprotein of ATPase activity at plasma membrane to efflux chemicals, addition of SRT2104 to the system was able to minimize such drug resistance acquired from senescent stromal cells.

Molecular profiles of sEVs produced via their biogenesis generally resemble those of their parental cells ^26^. Although the composition of sEVs encompasses multiple types of bioactive elements, here we selectively assessed the category of sRNA species particularly miRNA. This is basically due to the fact that the miRNA profile of sEVs can intimately mirror that of the parental donors correlating to a specific pathophysiological state but usually absent in healthy or normal cells, and these molecules have the potential to serve as possible biomarker for disease progression^62^. Here we found 53 miRNAs significantly upregulated in senescent stromal cells, although their relevance in cellular senescence remain largely unexplored. In addition to upregulated miRNAs, there are indeed downregulated miRNA such as miR-130b-3p, miR-214-3p and miR-7704, which can functionally target ABCB family members. However, which of these miRNAs is (or are) specifically linked with and mechanistically responsible for ABCB4 upregulation in recipient cancer cells upon uptake of senescent stromal sEVs, remains unclear yet. Further, contribution of each of these miRNAs to cancer malignancy including but not limited to acquired resistance, represents an intriguing topic and awaits future exploration.

There are increasing lines of evidence suggesting that sEVs may constitute part of the SASP by mediating paracrine effects on the nearby microenvironment ^63, 64^. A recent study using a Cre-reporter system demonstrated a positive correlation between sEV uptake and senescence activation, and identified IFITM3 within sEVs partially responsible for transmitting paracrine senescence to normal cells ^65^. Several studies have showed that cellular secretion of sEVs is enhanced in certain situations such as oxidative stress, telomeric attrition, irradiation and hypoxia, conditions that can induce cellular senescence ^62^. Although senescent cells (including those developed from cancer cells) release significantly more EVs than their non-senescent counterparts, and carry molecules involved in key pathophysiological processes ^65–69^, how stromal cells in the TME behave upon therapy-induced senescence (TIS) remains yet underexplored. Our data support that senescent stromal cells produce increased numbers of sEVs, which can be conceptually considered as part of the full SASP spectrum. In the TME niche, importantly, uptake of senescent stromal sEVs by recipient cancer cells markedly enhances their malignancy, including but not limited to acquired resistance. Bioactive contents such as miRNAs in these sEVs may serve as novel prognostic markers for evaluation of treatment consequence in cancer clinics. As a special note, combination of conventional chemotherapy, which was theoretically designed to target cancer cells, with a SIRT1 activator like SRT2104, can significantly improve therapeutic outcome by restraining the biogenesis of sEVs in senescent stromal cells, which frequently arise upon tissue damage triggered as a part of side-effects of anticancer agents and are responsible for multiple aging-related pathologies including but not limited to the most lethal form, cancer.

## MATERIALS AND METHODS

### Cell culture

Primary normal human prostate stromal cell line PSC27 and breast stromal cell line HBF1203 were maintained in stromal complete medium as described ^24^. Prostate epithelial cell lines PC3, DU145, LNCaP and breast epithelial cell line MDA-MB-231 (ATCC) were routinely cultured with RPMI1640 (10% FBS). The neoplastic and metastatic M12 was derived from BPH1, a benign prostate hyperplasia cell line immortalized by SV40-LT ^70, 71^. All cell lines were tested negative for mycoplasma contamination and authenticated with STR assays.

### *In vitro* treatments and senescence appraisal

PSC27 cells were grown until 80% confluent (CTRL) and treated with bleomycin (50 μg/ml, BLEO) for 12 h. After treatment, the cells were rinsed thrice with PBS and allowed to stay for 7-10 d in media. Alternatively, HBF1203 cells were treated with doxorubicin (10 μM, DOX) for 12 h to induce senescence. To examine cellular senescence, SA-β-Gal staining and BrdU incorporation were performed with the commercial kits (Beyotime and BD Biosciences, respectively) as previously described ^9, 57^. DNA-damage extent was evaluated by immunostaining for γH2AX or p-53BP1 foci by following a 4-category counting strategy as formerly reported ^24^. Random fields were chosen to show SA-β-gal positivity, BrdU incorporation or DDR foci, and quantified using CellProfiler (http://www.cellprofiler.org). For clonogenic assays, cells were seeded at 1 × 10^3^ cells/dish in 10 mm dishes for 24 h before treated with chemicals. Cells were fixed with 2% paraformaldehyde 7-10 d post treatment, gently washed with PBS and stained instantly with 10% crystal violet prepared in 50% methanol. Excess dye was removed with PBS, with plates photographed. Colony formation were evaluated by quantifying the number of single colonies per dish.

### Cancer patient recruitment and biospecimen analysis

Chemotherapeutic administration involving genotoxic agents was performed for primary prostate cancer patients (Clinical trial no. NCT03258320), by following the CONSORT 2010 Statement (updated guidelines for reporting parallel group randomized trials). Patients with a clinical stage ≥ I subtype A (IA) (T1a, N0, M0) of primary prostate cancer but without manifest distant metastasis were enrolled into the multicentered, randomized, double-blinded and controlled pilot studies. Patients with the age between 40-75 years with histologically proven prostate cancer were recruited into the clinical cohorts. Data regarding tumor size, histologic type, tumor penetration, lymph node metastasis, and TNM stage were obtained from the pathologic records. Before chemotherapy, tumors were acquired from these patients as ‘Pre’ samples (an ‘Untreated’ cohort). After chemotherapy, remaining tumors in patients were acquired as ‘Post’ samples (a ‘Chemo-treated’ cohort, with most tumors collected within 1-6 months after treatment). Tumors were processed as formalin-fixed paraffin-embedded (FFPE) biospecimens and sectioned for histological assessment, with alternatively prepared OCT-frozen chunks processed via lase capture microdissection (LCM) for gene expression analysis. For immunohistochemical (IHC) staining outputs, the immunoreactive scoring (IRS) gives a range of 1–4 qualitative scores according to staining intensity per tissue sample. Categories for the IRS include 0-1 (negative), 1-2 (weak), 2-3 (moderate), 3-4 (strong) ^72^. Tissue diagnosis was confirmed based on histological evaluation by independent pathologists. Randomized control trial (RCT) protocols and all experimental procedures were approved by the Ethics Committee and Institutional Review Board of Shanghai Jiao Tong University School of Medicine and Zhongshan Hospital of Fudan University, with methods carried out in accordance with the official guidelines. Informed consent was obtained from all subjects and the experiments conformed to the principles set out in the WMA Declaration of Helsinki and the Department of Health and Human Services Belmont Report.

### Cell phenotypic assays

Conditioned media (CM) from primary human stromal cells by brief centrifugation at 1500 g for 10 min at 4°C. A 6-well plate with cancer cells seeded at a density of 20,000/well was used for proliferation assays. For migration assays, a 1-ml micropipette tip was used to create linear scratches along the cell monolayer to simulate a wound at the vessel surface, and cell debris was removed by washing the cells once with basal medium. Images of each wound were taken at specific reference points along the scratch at designated timepoints. Image-Pro software was used to measure the total wound area to calculate percentage change in wound closure. For invasion assays, transwell inserts (8.0 μm pore size) (BD, catalog no. 353097) containing cancer cells at a cell density of 20,000/insert were fitted into the 6-well plate containing CM of stromal cells. The co-culture was incubated at 37°C and 5% CO2 for 3 d, with cells that have invaded through the transwell stained with DAPI and imaged with immunofluorescent microscopy. For chemoresistance assays, cancer cells were incubated with stromal cell-derived EVs, or with a chemotherapeutic agent MIT (or DOX) provided concurrently with stromal EVs in wells for 3 days at each cell line’s IC50, a value experimentally predetermined. The number of surviving cancer cells was normalized to the untreated group and presented as percentage of viability. For SIRT1 activation, compounds SRT1720 and SRT2104 were applied at the concentration of 2.5 μM each case and used alone or together with genotoxic agents. For SIRT1 inhibition, agents SAHA and NAM were used at the concentration of 10 nM and 200 nM, respectively, either alone for combined with genotoxic drugs to treat stromal cells.

### EV preparation

The bovine EV-depleted media were obtained by overnight ultracentrifugation at 100,000 × g in a 45Ti rotor (Beckman Coulter), of DMEM supplemented with 10% FBS. On day 7 or 8 after stromal cell treatment in culture or when subconfluence was reached in the proliferating group (control), cells were washed in PBS and further cultured in EV-depleted medium for 24 h before collection of CM for EV isolation. EVs were enriched by differential ultracentrifugation as previously reported ^73^. Briefly, CM was centrifuged at 300 × g for 10 min at 4 °C to pellet cells. Supernatant was centrifuged at 2,000 × g for 20 min at 4 °C to remove cells and cell debris (2K pellet), transferred to new tubes, and centrifuged in a 45Ti rotor for 40 min at 10,000 × g (9,000 rpm = 10K pellet), and finally for 90 min at 100,000 × g (30,000 rpm = 100K) in a P55ST2-639 rotor with an Avanti JXN-26 (Beckman) centrifuge to collect EV pellet. Final pellets were washed in 50-60 ml of PBS and re-centrifuged at 100,000 × g to eliminate contaminant proteins and nucleic acids, before resuspended in 50-100 μl of sterile PBS then stored at −80°C until use. Cells recovered from the first 300 × g pellet were pooled with cells detached from the culture plates by incubation at 4 °C in trypsin-EDTA (Gibco) as adherent cells and counted by Countess (Invitrogen). Viability was assessed by Trypan Blue stain 0.4% (Life Technologies) exclusion.

### Iodixanol/Optiprep gradient separation

For iodixanol-based gradient separation, pellets obtained by ultracentrifugation from 80 to 200 million of stroma cells were washed and resuspended in 1.5 ml buffer containing: 0.25 M sucrose, 10 mM Tris pH 8.0, 1 mM EDTA (pH 7.4), transferred to a SW55Ti rotor tube (Beckman), and mixed 1:1 with 60% (wt/vol) stock solution of iodixanol/Optiprep. A 40% iodixanol working solution was prepared [40% (wt/vol) iodixanol, 0.25 M sucrose, 10 mM Tris pH 8.0, 1 mM EDTA, final pH set to 7.4] and used to prepare 20% and 10% (wt/vol) iodixanol solutions. Next, 1.3 ml 20% (wt/vol) iodixanol and 1.2 ml 10% iodixanol were successively layered on top of the vesicle suspension and tubes were centrifuged for 1 h at 4 °C at 350,000 × g (54,000 rpm) in SW55Ti (stopping without break, DECEL=0); 10 fractions of 490 μl were collected from the top of the tube as described formerly ^55^. Density was assessed with a refractometer (Carl Zeiss). Fractions were diluted with 2.5 ml sterile PBS and centrifuged for 30 min at 100,000 × g (43,000 rpm) in a TLA 110 rotor (Beckman, Optima TL100 centrifuge). These concentrated fractions were resuspended in 30 μl sterile PBS prior to further assessment by electron microscopy or RNA preparation for sequencing.

### Nanoparticle tracking analysis (NTA)

EVs were subject to characterization by the NTA technique. EVs isolated and resuspended in PBS were analyzed for size and concentration via NTA using a NanoSight NS300 device (Malvern Panalytical) equipped with a 532-nm laser, low-volume flow-cell, syringe pump, and concentration upgrade. Thawed EV samples were further diluted in PBS to yield concentrations of between 20 and 100 particles/frame on NanoSight, and at least five 30-60s videos were acquired per sample using the same camera settings. At least one dilution, videos were taken and analyzed by the NTA 3.1 or 3.2 software with default settings.

For purification of sEVs and MVs, fresh cell culture supernatant was subject to successive differential ultracentrifugation [30]. Briefly, after two steps of 300 × g for 10 min at 4 °C to pellet cells and debris, the supernatant (SN) was centrifuged at 14,000 × g for 35 min, with the resultant pellet (P14) washed in PBS. SN14 was centrifuged at 100,000 × g for 2 h, with the pellet (P100) washed in PBS. EV concentration in SN0.3 or P14/P100 were generally normalized to the cell number. For instance, the supernatant of 1 × 10^6^ cells was used for EV purification and concentration of P14 and P100 (resuspended in 100 μl PBS and stored at −20°C) was measured by NTA. Before assays, P14 and P100 samples were diluted between 8- and 100-fold in PBS to obtain the optimal measurement concentrations of 5-15 ×10^8^ particles/ml, with EV concentration calculated by using 10 bins of 50 nm each until 90% of all EVs were covered.

### Electron microscopy

Stromal cells or their derived EVs were precisely imaged by transmission electron microscopy (TEM). Electron microscopy was performed on pellets stored at −80 °C and never unfrozen. Cell or EV suspension in PBS was deposited on formvar-carbon–coated cooper/palladium grids for whole-mount analysis as described previously ^55, 74^. In brief, cells or EVs were incubated overnight on carbon film 400 square mesh TEM grids. Grids were briefly dipped in a droplet of ultrapure water, wicked dry, vacuum dried, and imaged on a Philips CM120 (Philips Research).

### Stromal cell lysosomal pH assay

Lysosomal pH was measured as reported previously ^75^, with minor modifications. Briefly, cells were treated with 50 μg/ml of LysoSensor Yellow/Blue Dextran (10,000 MW) for 12 h, before amino acid-starved for an additional 2 h. The cells were then rinsed twice with PBS, and resuspended in physiological buffer (136 mM NaCl, 2.5 mM KCl, 2 mM CaCl_2_, 1.3 mM MgCl_2_, 5mM Glucose, 10mM HEPES, pH 7.4), and transferred to individual wells of a black 96-well plate. LysoSensor fluorescence emission was recorded at 460 nm and 540 nm upon excitation at 360 nm using a BioTek Synergy II Plate Reader. To measure the lysosomal pH, the 460/540 fluorescence emission ratios were interpolated to a calibration curve that was established by resuspending the cells containing lysoSensor in 200 μl aliquots of pH calibration buffers (145 mM KCl, 10 mM glucose, 1 mM MgCl2, and 20 mM of either HEPES, MES, or acetate supplemented with 10 μg/ml nigericin), buffered to pH ranging from 3.5 to 8.0.

### Lysosomal pH recovery assay

Lysosomal pH was detected as described previously ^76^, with minor modifications. Cells (5 × 10^3^) plated within 24 well plates were treated with either DMSO, or 100 nM Bafilomycin-A, for 1 h, at which point the cells were rinsed extensively with media and allowed to recover for 1 h. The cells were then treated with LysoTracker Green DND-26 (Thermo Fisher) for 1 h, washed twice with PBS, and lysed in 200 μl of RIPA buffer (10 mM Tris, 140 mM NaCl, 1% Triton X-100, 1 mM EDTA, 0.1% SDS, 0.1% sodium deoxycholate and 1 μg/ml each aprotinin and leupeptin). The fluorescence was measured in a 96-well plate using a BioTek Synergy II Plate Reader; excitation at 485 nm and emission at 520 nm, respectively. The percent recovery was calculated as follows: 100%: DMSO for the whole duration of the assay (*A_100_*); 0%: Bafilomycin-A for the whole duration of the assay% (*A_0_*); Recovery = (*A_R_*−*A*_0_)/(*A*_100_−*A*_0_).

### Library preparation, RNA-seq and bioinformatics analysis

Total RNA was extracted from human stromal cells using Trizol reagent according the Manufacturer’s instructions (Invitrogen) and genomic DNA was removed using rDNase I RNase-free (Takara). RNA quality was verified by both ND-2000 (NanoDrop) and Bioanalyzer 2100 (Agilent), with only high-quality RNA sample (OD260/280 = 1.8∼2.2, OD260/230 ≥ 2.0, RIN ≥ 8, 28S:18S ≥ 1.0) taken. Totally 3 μg of total RNA was ligated with 5′- and a 3′-adaptors sequentially with TruSeq^TM^ Small RNA (or Stranded Total RNA in the case of cancer cells) Sample Prep Kit (Illumina). Briefly, cDNA was synthesized by reverse transcription and amplified with 12 PCR cycles to produce libraries. After quantified by TBS380, deep sequencing was performed on the Illumina HiSeq X10 sequencing platform by Shanghai Majorbio Bio-Pharm Biotechnology Co., Ltd. (Shanghai, China).

Paired-end transcriptomic reads were mapped to the reference genome (GRCh38/hg38) with reference annotation from Gencode v27 using the Bowtie tool. Duplicate reads were identified using the picard tools (1.98) script mark duplicates (https://github.com/broadinstitute/picard) and only non-duplicate reads were retained. Reference splice junctions are provided by a reference transcriptome (Ensembl build 73) ^77^. FPKM values were calculated using Cufflinks, with differential gene expression called by the Cuffdiff maximum-likelihood estimate function ^78^. Genes of significantly changed expression were defined by a false discovery rate (FDR)-corrected *P* value < 0.05. Only ensembl genes 73 of status “known” and biotype “coding” were used for downstream analysis.

Reads were trimmed using Trim Galore (v0.3.0) (http://www.bioinformatics.babraham.ac.uk/projects/trim_galore/) and quality assessed using FastQC (v0.10.0) (http://www.bioinformatics.bbsrc.ac.uk/projects/fastqc/). Differentially expressed genes were subsequently analyzed for enrichment of biological themes using the DAVID bioinformatics platform (https://david.ncifcrf.gov/), the Ingenuity Pathways Analysis program (http://www.ingenuity.com/index.html). Raw data were preliminarily analyzed on the free online platform of Majorbio I-Sanger Cloud Platform (www.i-sanger.com), and subsequently deposited in the NCBI Gene Expression Omnibus (GEO) database.

#### Venn diagrams

Venn diagrams and associated empirical *P*-values were generated using the USeq (v7.1.2) tool IntersectLists ^79^. The t-value used was 22,008, as the total number of genes of status “known” and biotype “coding” in ensembl genes 73. The number of iterations used was 1,000.

#### RNA-seq heatmaps

For each gene, the FPKM value was calculated based on aligned reads, using Cufflinks ^78^. Z-scores were generated from FPKMs. Hierarchical clustering was performed using the R package heatmap.2 and the distfun = “pearson” and hclustfun = “average”.

#### Protein-protein interaction network

Protein-protein interaction (PPI) analysis was performed with STRING 3.0 ^47^. The specific proteins meeting the criteria, were imported to NetworkAnalyst (http://www.networkanalyst.ca) ^80^. A minimum interaction network was chosen for further hub and module analysis.

### Plasma isolation and ELISA assays

The peripheral blood of prostate cancer patients was acquired from Zhongshan Hospital of Fudan University, following an IRB-approved sample collection protocol. Clinical investigation procedures were compliant with all relevant ethical regulations regarding human subjects, who provided signed consensus for venous blood collection. Serum was isolated by centrifugation of raw plasma and subject to a TSG101-specific self-coated ELISA assay.

### Preclinical studies

All animals were maintained in a specific pathogen-free (SPF) facility, with NOD/SCID (Nanjing Biomedical Research Institute of Nanjing University) mice at an age of approximately 6 weeks (∼20g body weight) used. Each group comprised 10 mice, and xenografts were subcutaneously implanted at the hind flank. Stromal cells (PSC27) were mixed with cancer cells (PC3) at a ratio of 1:4 (250,000 stromal cells admixed with 1,000,000 cancer cells to make tissue recombinants). Animals were sacrificed at end of the 8th week post tumor xenografting, with tumor volume measurement, tissue staining and histological examination performed as previously described ^23^.

For chemoresistance studies, animals received subcutaneous implantation of tissue recombinants as described above and were given standard laboratory diets for 2 weeks to allow tumor uptake and growth initiation. Starting from the 3^rd^ week (tumors reaching 4-8 mm in diameter), MIT (0.2 mg/kg doses) or vehicle controls was administered by body injection (chemicals via intraperitoneal route), on the 1^st^ day of 3^rd^, 5^th^ and 7^th^ weeks, respectively. Upon completion of the 8-week regimen, animals were sacrificed, with tumor volumes recorded and tissues processed for histological evaluation. For SIRT1 activation, SRT2104 (Selleck) was dissolved in suspension (30% PEG 400/0.5% Tween 80/5% Propylene glycol at 30 mg/ml), administered via oral gavage at a dose of 100 mg/kg bodyweight every other week, starting from the 1st day of the 3^rd^ week after xenograft implantation and administered either alone or together with MIT, until completion of the 8-week regimen.

All animal experiments were performed in compliance with NIH Guide for the Care and Use of Laboratory Animals (National Academies Press, 2011) and the ARRIVE guidelines, and were approved by the Institutional Animal Care and Use Committee (IACUC) of Shanghai Institutes for Biological Sciences, Chinese Academy of Sciences.

### Statistical analyses

All *in vitro* experiments were performed in triplicates, and animal studies were conducted with n ≥ 8 mice per group. Animals were distributed into groups of equal body weight, and no animals were excluded from analysis. Sample sizes were estimated based on an 80% power to detect a 50% reduction in tumor volumes observed in mice subject to chemotherapy and/or targeted therapy compared with control mice, accepting a type I error rate of 0.05.

When comparing two conditions, Student’s *t* tests with Mann-Whitney’s tests were used to determine statistical significance. More than two comparisons were made using one-way ANOVA followed by Dunn’s or Tukey’s post hoc tests. Two-way ANOVA with Bonferroni correction was used for grouped analysis. Nonlinear dose-response fitting curves were generated with GraphPad Prism 7.0. *P* values < 0.05 were considered significant, and two-sided tests were performed. Unless otherwise indicated, all data in the figures were presented as mean ± SD. Univariate and multivariate Cox proportional hazards model analysis were performed with statistical software SPSS.

### Data availability

The raw RNA-seq data have been deposited in the Gene Expression Omnibus database (accession code GSE128282, GSE128927 and GSE128405). The authors declare that all other data supporting the findings of this study are available within the article or its Supplementary Information and from the corresponding author upon reasonable request.

## ACKNOWLEDGEMENTS

This work was supported by grants from National Key Research and Development Program of China (2016YFC1302400), National Natural Science Foundation of China (NSFC) (81472709, 31671425, 31871380) to Y.S., CAS Key Laboratory of Tissue Microenvironment and Tumor of Chinese Academy of Sciences; National Science and Technology Major Project of China (2019ZX09732002-14) to J.J.; Breast Cancer Now (2012MayPR070; 2012NovPhD016), the Medical Research Council of the United Kingdom (MR/N012097/1), Cancer Research UK Imperial Centre, Imperial ECMC and NIHR Imperial BRC to E.W-F.L.

## AUTHOR CONTRIBUTIONS

Y.S. conceived this study, designed the experiments and supervised the project. L.H. carried out most of the biological experiments. Q.L. acquired and analyzed clinical samples from prostate cancer patients, and managed subject information. L.H. and Y.S. performed data mining and bioinformatics of gene expression and signaling pathways. S.L., Q.X., B.Z., X.D., M.Q. and Y.J. helped *in vitro* culture and phenotypic characterization of cancer cells. J.G, L.C., Y.E.C. E.W-F.L. and J.J. provided conceptual inputs or supervised a specific subset of experiments. L.H., Q.L. and Y.S. performed preclinical studies. Y.S. prepared the manuscript. All authors critically read and commented on the final manuscript.

## ADDITIONAL INFORMATION

Supplementary information accompanies this paper online at Cell Research website.

