## Supplementary Information for "Senescent stromal cells promote cancer resistance through SIRT1 loss-potentiated overproduction of small extracellular vesicles"

**This PDF file includes:**

Supplementary Information Fig. S1 Cellular senescence, expression profiling and the SASP development in human stromal cells upon genotoxic treatment.
Supplementary Information Fig. S2 Statistical analysis of miRNAs differentially carried by sEVs of PRE vs SEN stromal cells and database mapping of their targets in human.
Supplementary Information Fig. S3 SIRT1 decline is not related with single strand breaks, proteasome degradation or autophagy deficiency upon cellular senescence.
Supplementary Information Fig. S4 Senescent stromal sEVs enhance the malignancy of prostate cancer cells.
Supplementary Information Fig. S5 Transcriptomic profiling of prostate cancer cells upon exposure to senescent stromal sEVs and establishment of protein-protein interaction network involving ABCB4.
Supplementary Information Fig. S6 Senescent stromal sEV-conferred malignancy is diminished upon elimination of ABCB4 from cancer cells.
Supplementary Information Fig. S7 Pharmacological targeting SIRT1 minimizes senescent stromal sEV production and prevents ABCB4 upregulation in prostate cancer cells.

Supplementary Information Table S1 Statistics of small RNA species upon global assessment of Illumina sequencing data

Supplementary Information Table S2 Representative list of the molecular targets of upregulated and downregulated miRNAs in SEN stromal cell-derived sEVs

Supplementary Information Table S3 Predicted miRNA target molecules located at the key nodes of intracellular protein-protein interaction (PPI) network

Supplementary Information Table S4 Functional enrichments of predicted partners of ABCB4 in PPI network

Supplementary Information Table S5 Primer sequences for qRT-PCR assays

Supplementary Information Table S6 Antibodies used for immunoblot, immunofluorescence and immunohistochemistry staining

### Figure S1

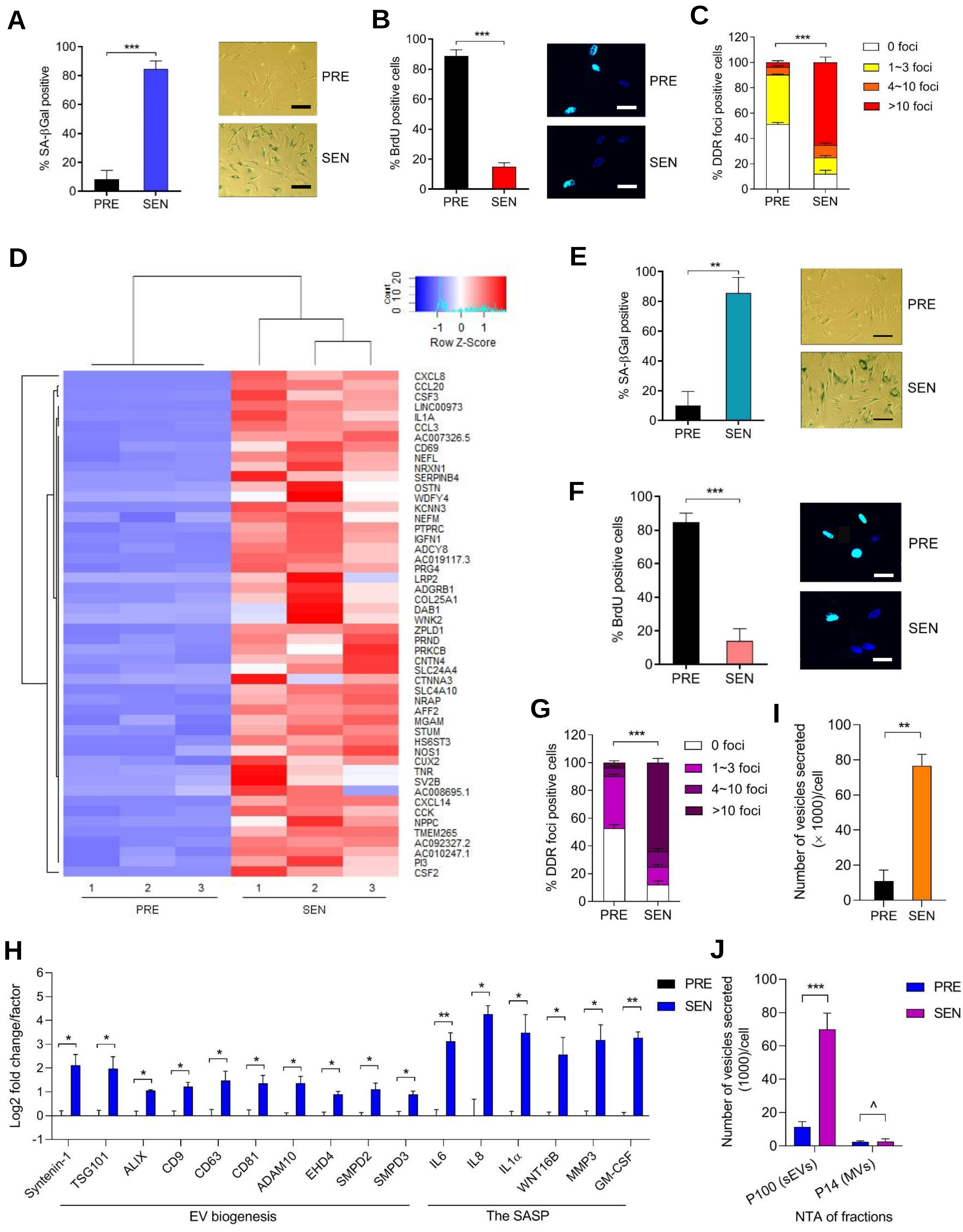
A
% SA-βGal positive
***
PRE
SEN
B
% BrdU positive cells
C
% DDR foci positive cells
0 foci
1~3 foci
4~10 foci
>10 foci
D
Row Z-Score
Count
CXCL8
CCL20
CSF3
LINC00973
IL1A
CCL3
AC007326.5
CD69
NEFL
NRXN1
SERPINB4
OSTN
WDFY4
KCNN3
NEFM
PTPRC
IGFN1
ADCY8
AC019117.3
PRG4
LRP2
ADGRB1
COL25A1
DAB1
WNK2
ZPLD1
PRND
PRKCB
CNTN4
SLC24A4
CTNNA3
SLC4A10
NRAP
AFF2
MGAM
STUM
HS6ST3
NOS1
CUX2
TNR
SV2B
AC008695.1
CXCL14
CCK
NPPC
TMEM265
AC092327.2
AC010247.1
PI3
CSF2
E
F
G
I
Number of vesicles secreted (× 1000)/cell
H
Log2 fold change/factor
Syntenin-1
TSG101
ALIX
CD9
CD63
CD81
ADAM10
EHD4
SMPD2
SMPD3
IL6
IL8
IL1α
WNT16B
MMP3
GM-CSF
EV biogenesis
The SASP
J
Number of vesicles secreted (1000)/cell
P100 (sEVs)
P14 (MVs)
NTA of fractions

### Figure S2

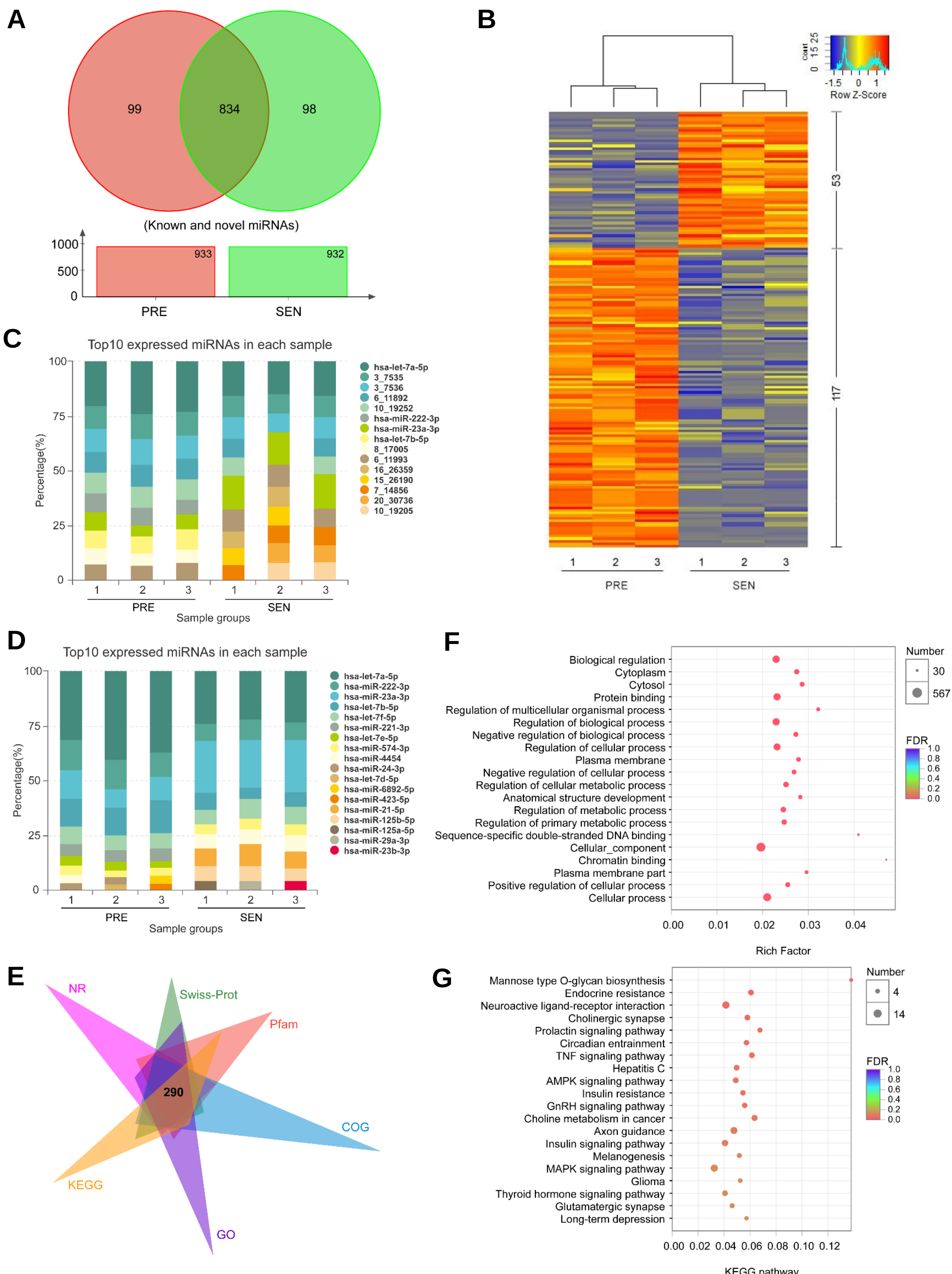
A
99
834
98
(Known and novel miRNAs)
1000
500
0
933
932
PRE
SEN
B
Count
Row Z-Score
53
117
1 2 3 1 2 3
PRE
SEN
C
Top10 expressed miRNAs in each sample
Percentage(%)
Sample groups
hsa-let-7a-5p
3_7535
3_7536
6_11892
10_19252
hsa-miR-222-3p
hsa-miR-23a-3p
hsa-let-7b-5p
8_17005
6_11993
16_26359
15_26190
7_14856
20_30736
10_19205
D
Top10 expressed miRNAs in each sample
Percentage(%)
Sample groups
hsa-let-7a-5p
hsa-miR-222-3p
hsa-miR-23a-3p
hsa-let-7b-5p
hsa-let-7f-5p
hsa-miR-221-3p
hsa-let-7e-5p
hsa-miR-574-3p
hsa-miR-4454
hsa-miR-24-3p
hsa-let-7d-5p
hsa-miR-6892-5p
hsa-miR-423-5p
hsa-miR-21-5p
hsa-miR-125b-5p
hsa-miR-125a-5p
hsa-miR-29a-3p
hsa-miR-23b-3p
E
NR
Swiss-Prot
Pfam
COG
KEGG
GO
290
F
Biological regulation
Cytoplasm
Cytosol
Protein binding
Regulation of multicellular organismal process
Regulation of biological process
Negative regulation of biological process
Regulation of cellular process
Plasma membrane
Negative regulation of cellular process
Regulation of cellular metabolic process
Anatomical structure development
Regulation of metabolic process
Regulation of primary metabolic process
Sequence-specific double-stranded DNA binding
Cellular_component
Chromatin binding
Plasma membrane part
Positive regulation of cellular process
Cellular process
Number
30
567
FDR
Rich Factor
G
Mannose type O-glycan biosynthesis
Endocrine resistance
Neuroactive ligand-receptor interaction
Cholinergic synapse
Prolactin signaling pathway
Circadian entrainment
TNF signaling pathway
Hepatitis C
AMPK signaling pathway
Insulin resistance
GnRH signaling pathway
Choline metabolism in cancer
Axon guidance
Insulin signaling pathway
Melanogenesis
MAPK signaling pathway
Glioma
Thyroid hormone signaling pathway
Glutamatergic synapse
Long-term depression
Number
4
14
FDR
KEGG pathway

### Figure S3

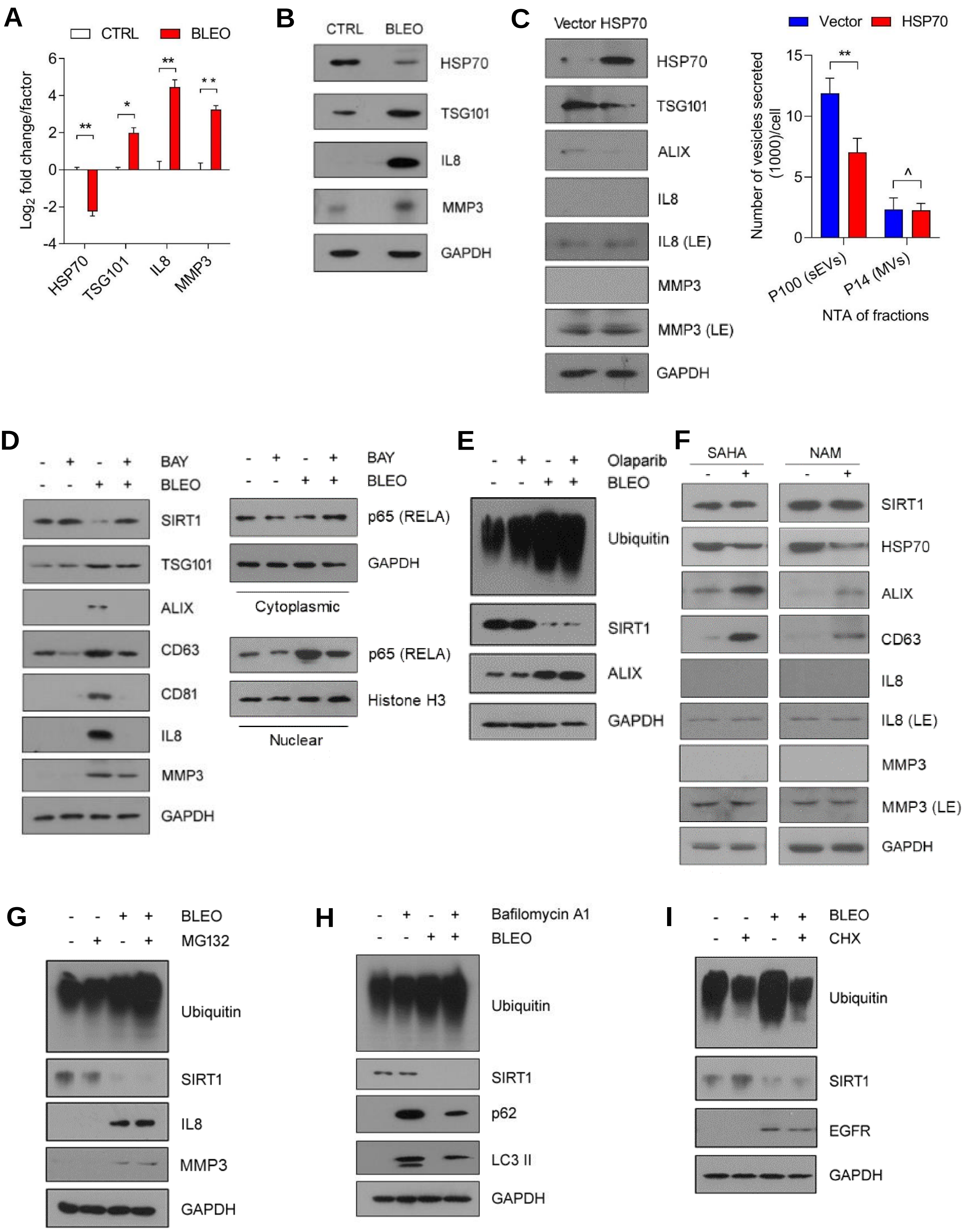
A
CTRL
BLEO
Log2 fold change/factor
HSP70
TSG101
IL8
MMP3
B
CTRL
BLEO
HSP70
TSG101
IL8
MMP3
GAPDH
C
Vector HSP70
HSP70
TSG101
ALIX
IL8
IL8 (LE)
MMP3
MMP3 (LE)
GAPDH
Vector
HSP70
Number of vesicles secreted (1000)/cell
P100 (sEVs)
P14 (MVs)
NTA of fractions
D
BAY
BLEO
SIRT1
TSG101
ALIX
CD63
CD81
IL8
MMP3
GAPDH
p65 (RELA)
GAPDH
Cytoplasmic
p65 (RELA)
Histone H3
Nuclear
E
Olaparib
BLEO
Ubiquitin
SIRT1
ALIX
GAPDH
F
SAHA
NAM
SIRT1
HSP70
ALIX
CD63
IL8
IL8 (LE)
MMP3
MMP3 (LE)
GAPDH
G
BLEO
MG132
Ubiquitin
SIRT1
IL8
MMP3
GAPDH
H
Bafilomycin A1
BLEO
Ubiquitin
SIRT1
p62
LC3 II
GAPDH
I
BLEO
CHX
Ubiquitin
SIRT1
EGFR
GAPDH

**Figure S4**

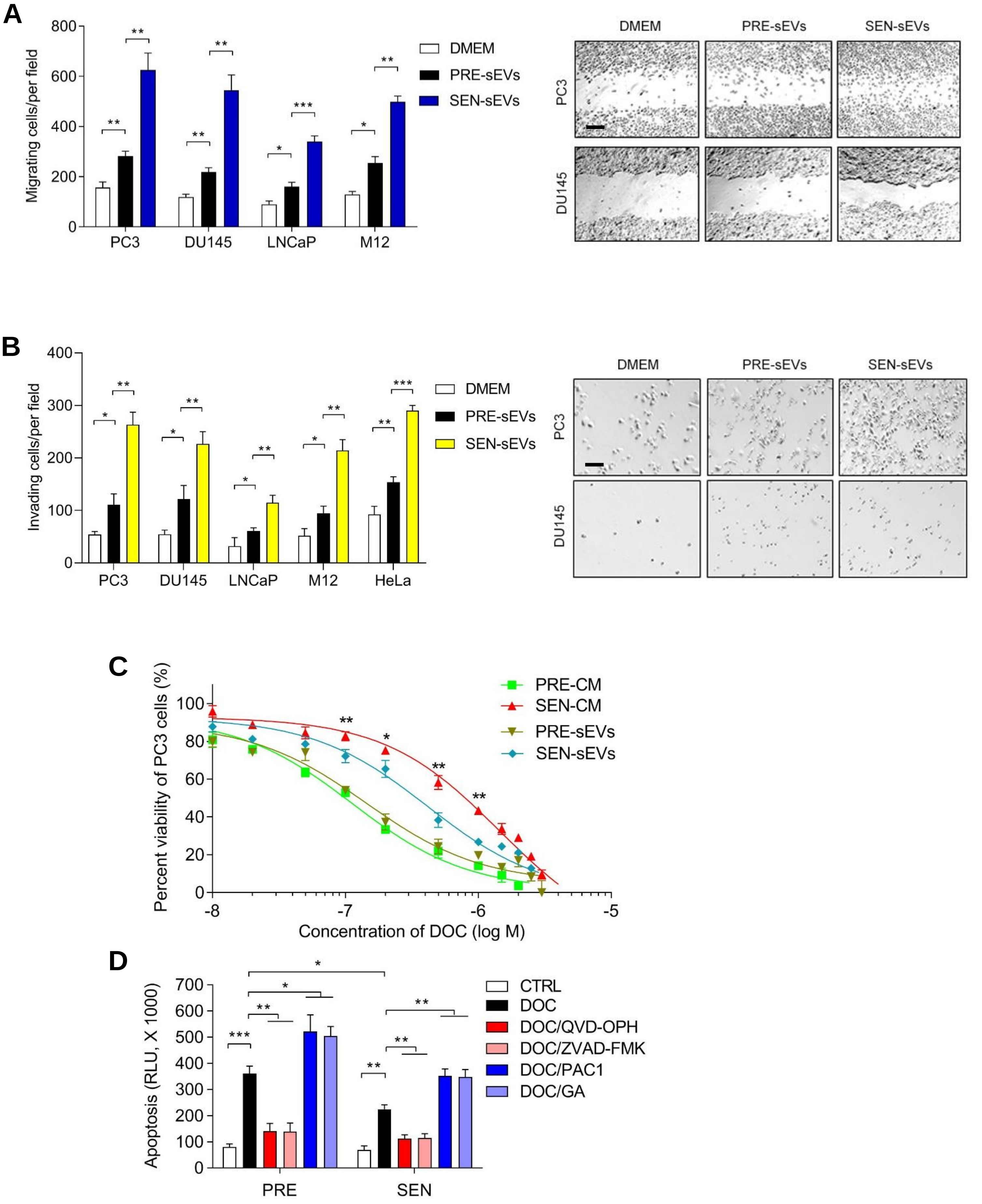
A
Migrating cells/per field
800
600
400
200
0
PC3
DU145
LNCaP
M12
DMEM
PRE-sEVs
SEN-sEVs
DMEM
PRE-sEVs
SEN-sEVs
PC3
DU145
B
Invading cells/per field
400
300
200
100
0
PC3
DU145
LNCaP
M12
HeLa
DMEM
PRE-sEVs
SEN-sEVs
DMEM
PRE-sEVs
SEN-sEVs
PC3
DU145
C
Percent viability of PC3 cells (%)
100
80
60
40
20
0
-8
-7
-6
-5
Concentration of DOC (log M)
PRE-CM
SEN-CM
PRE-sEVs
SEN-sEVs
D
Apoptosis (RLU, X 1000)
700
600
500
400
300
200
100
0
PRE
SEN
CTRL
DOC
DOC/QVD-OPH
DOC/ZVAD-FMK
DOC/PAC1
DOC/GA

**Figure S5**

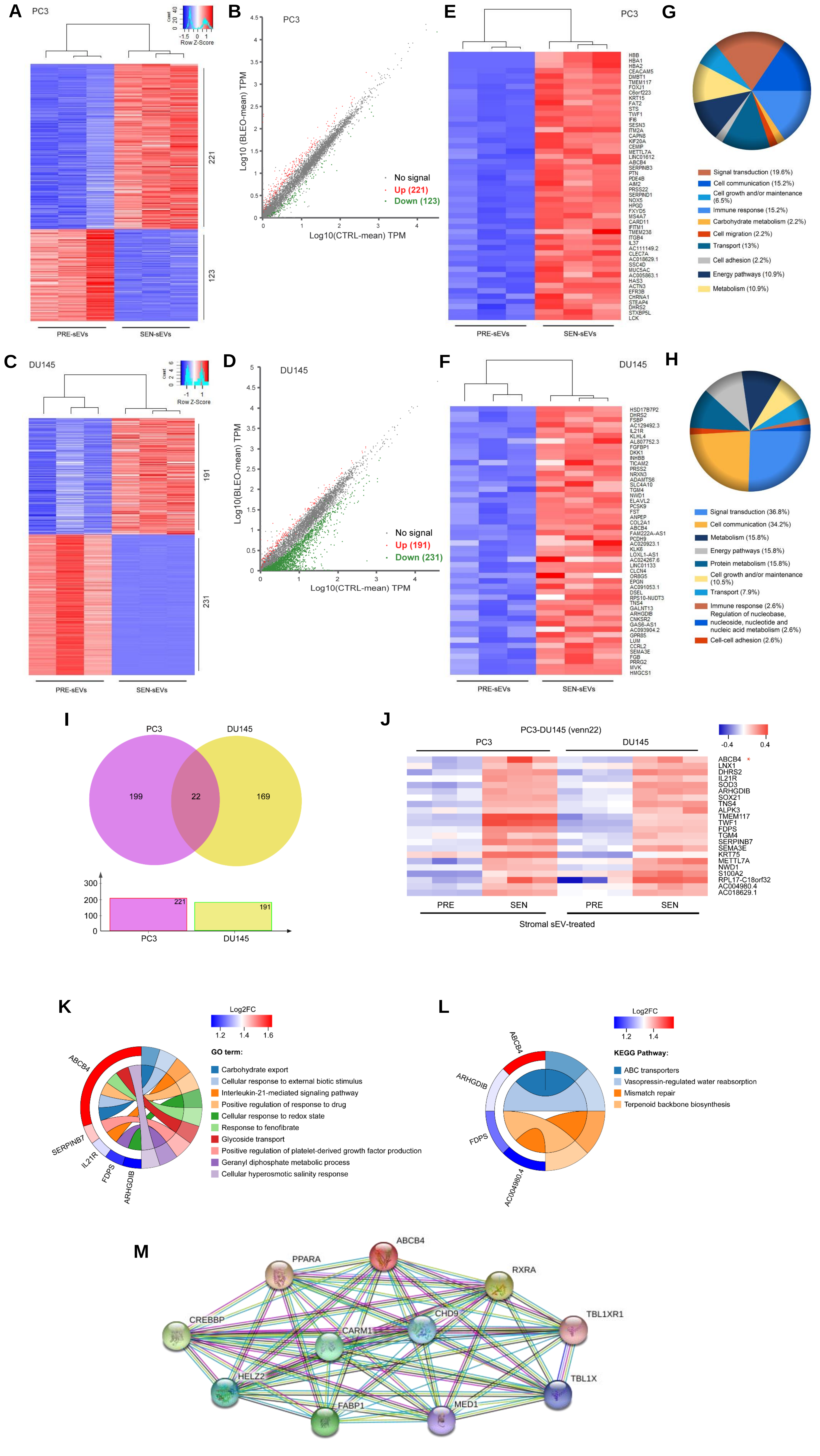

A
PC3
Count
Row Z-Score
221
123
PRE-sEVs
SEN-sEVs
B
PC3
Log10 (BLEO-mean) TPM
Log10(CTRL-mean) TPM
No signal
Up (221)
Down (123)
E
PC3
HBB
HBA1
HBA2
CEACAM5
DMBT1
TMEM117
FOXJ1
C6orf223
KRT15
FAT2
STS
TWF1
IFI6
SESN3
ITM2A
CAPN8
KIF20A
CEMIP
METTL7A
LINC01612
ABCB4
SERPINB3
PTN
PDE4B
AIM2
PRSS22
SERPIND1
NOX5
HPGD
FXYD5
MS4A7
CARD11
IFITM1
TMEM238
ITGB4
IL37
AC111149.2
CLEC7A
AC018629.1
SSC4D
MUC5AC
AC005863.1
HAS3
ACTN3
EFR3B
CHRNA1
STEAP4
DHRS2
STXBP5L
LCK
PRE-sEVs
SEN-sEVs
G
Signal transduction (19.6%)
Cell communication (15.2%)
Cell growth and/or maintenance (6.5%)
Immune response (15.2%)
Carbohydrate metabolism (2.2%)
Cell migration (2.2%)
Transport (13%)
Cell adhesion (2.2%)
Energy pathways (10.9%)
Metabolism (10.9%)
C
DU145
Count
Row Z-Score
191
231
PRE-sEVs
SEN-sEVs
D
DU145
Log10(BLEO-mean) TPM
Log10(CTRL-mean) TPM
No signal
Up (191)
Down (231)
F
DU145
HSD17B7P2
DHRS2
FSBP
AC129492.3
IL21R
KLHL4
AL807752.3
FGFBP1
DKK1
INHBB
TICAM2
PRSS2
NRXN3
ADAMTS6
SLC4A10
TGM4
NWD1
ELAVL2
PCSK9
FST
ANPEP
COL2A1
ABCB4
FAM222A-AS1
PCDH9
AC020923.1
KLK6
LOXL1-AS1
AC024267.6
LINC01133
CLCN4
OR8G5
EPGN
AC091053.1
DSEL
RPS10-NUDT3
TNS4
GALNT13
ARHGDIB
CNKSR2
GAS6-AS1
AC093904.2
GPR85
LUM
CCRL2
SEMA3E
FGB
PRRG2
MVK
HMGCS1
PRE-sEVs
SEN-sEVs
H
Signal transduction (36.8%)
Cell communication (34.2%)
Metabolism (15.8%)
Energy pathways (15.8%)
Protein metabolism (15.8%)
Cell growth and/or maintenance (10.5%)
Transport (7.9%)
Immune response (2.6%)
Regulation of nucleobase, nucleoside, nucleotide and nucleic acid metabolism (2.6%)
Cell-cell adhesion (2.6%)
I
PC3
DU145
199
22
169
221
191
PC3
DU145
J
PC3-DU145 (venn22)
PC3
DU145
-0.4
0.4
ABCB4 *
LNX1
DHRS2
IL21R
SOD3
ARHGDIB
SOX21
TNS4
ALPK3
TMEM117
TWF1
FDPS
TGM4
SERPINB7
SEMA3E
KRT75
METTL7A
NWD1
S100A2
RPL17-C18orf32
AC004980.4
AC018629.1
PRE
SEN
PRE
SEN
Stromal sEV-treated
K
Log2FC
GO term:
Carbohydrate export
Cellular response to external biotic stimulus
Interleukin-21-mediated signaling pathway
Positive regulation of response to drug
Cellular response to redox state
Response to fenofibrate
Glycoside transport
Positive regulation of platelet-derived growth factor production
Geranyl diphosphate metabolic process
Cellular hyperosmotic salinity response
ABCB4
SERPINB7
IL21R
FDPS
ARHGDIB
L
Log2FC
KEGG Pathway:
ABC transporters
Vasopressin-regulated water reabsorption
Mismatch repair
Terpenoid backbone biosynthesis
ABCB4
ARHGDIB
FDPS
AC004980.4
M
ABCB4
PPARA
RXRA
CREBBP
CHD9
TBL1XR1
CARM1
HELZ2
TBL1X
FABP1
MED1

### Figure S6

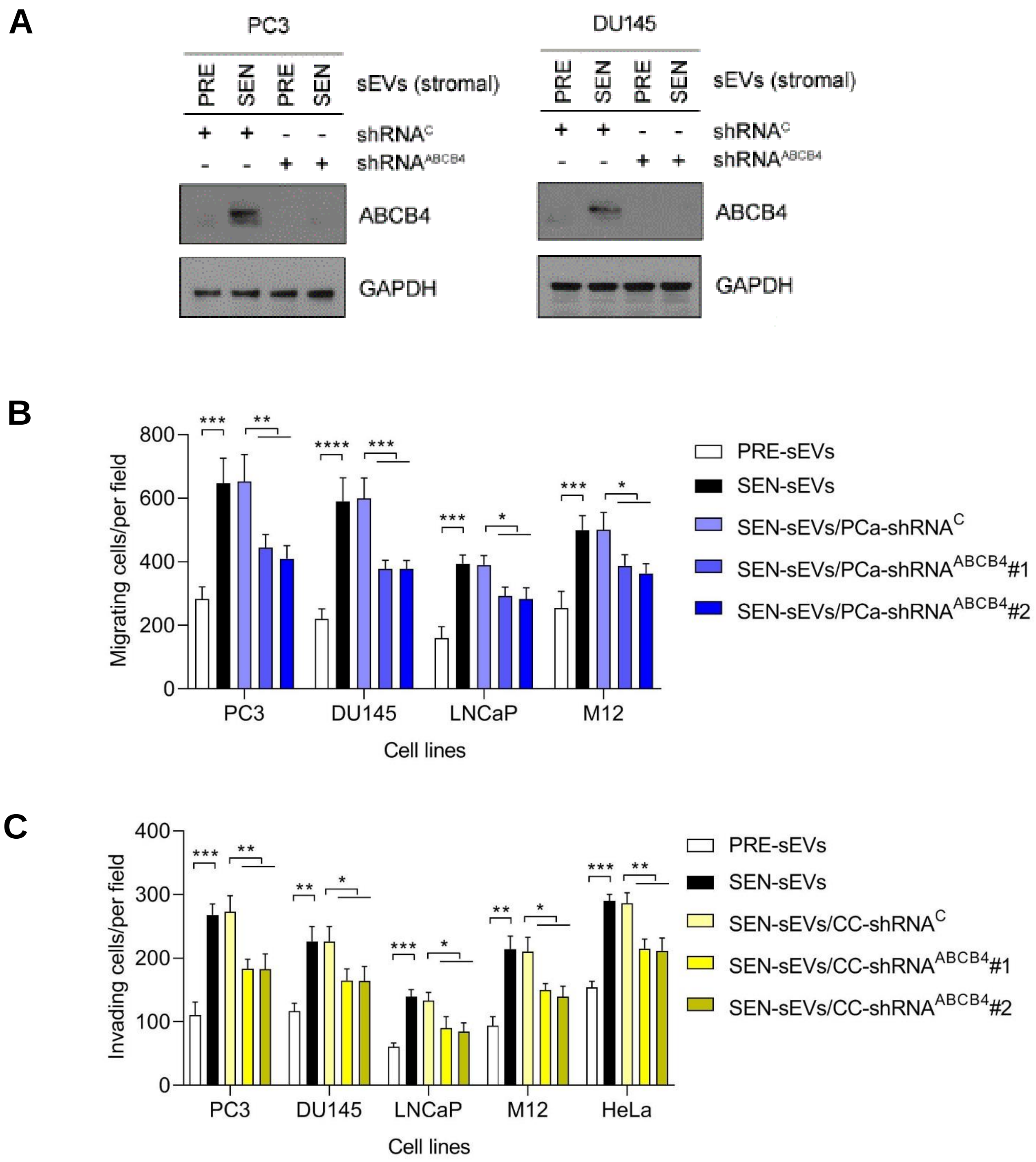
A
PC3
DU145
PRE SEN PRE SEN
sEVs (stromal)
shRNA^C
shRNA^ABCB4
ABCB4
GAPDH
B
Migrating cells/per field
800
600
400
200
0
PC3
DU145
LNCaP
M12
Cell lines
PRE-sEVs
SEN-sEVs
SEN-sEVs/PCa-shRNA^C
SEN-sEVs/PCa-shRNA^ABCB4#1
SEN-sEVs/PCa-shRNA^ABCB4#2
C
Invading cells/per field
400
300
200
100
0
PC3
DU145
LNCaP
M12
HeLa
Cell lines
PRE-sEVs
SEN-sEVs
SEN-sEVs/CC-shRNA^C
SEN-sEVs/CC-shRNA^ABCB4#1
SEN-sEVs/CC-shRNA^ABCB4#2

Figure S7

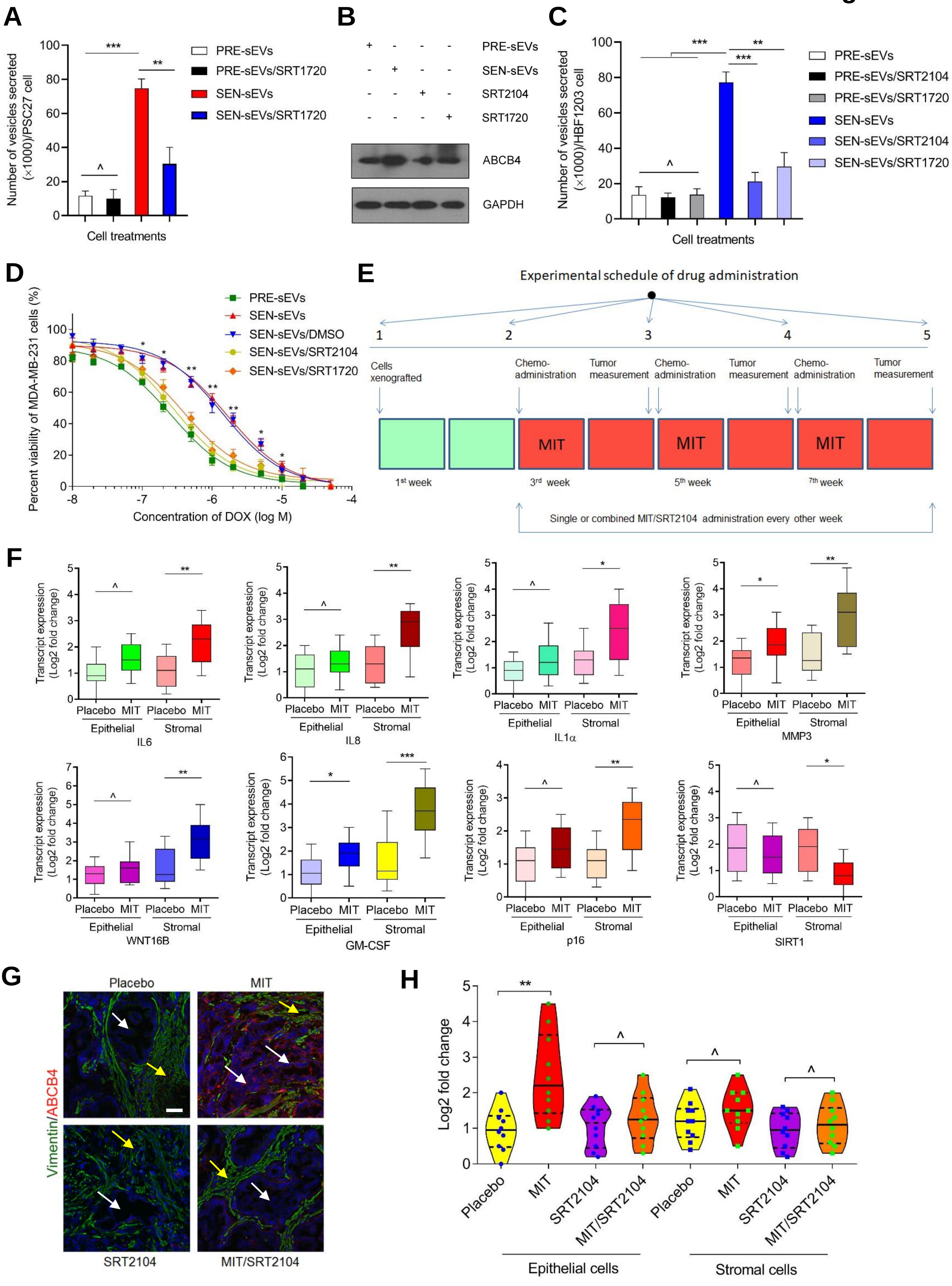
A
Number of vesicles secreted (×1000)/PSC27 cell
Cell treatments
PRE-sEVs
PRE-sEVs/SRT1720
SEN-sEVs
SEN-sEVs/SRT1720
***
**
^
B
PRE-sEVs
SEN-sEVs
SRT2104
SRT1720
ABCB4
GAPDH
C
Number of vesicles secreted (×1000)/HBF1203 cell
Cell treatments
PRE-sEVs
PRE-sEVs/SRT2104
PRE-sEVs/SRT1720
SEN-sEVs
SEN-sEVs/SRT2104
SEN-sEVs/SRT1720
D
Percent viability of MDA-MB-231 cells (%)
Concentration of DOX (log M)
PRE-sEVs
SEN-sEVs
SEN-sEVs/DMSO
SEN-sEVs/SRT2104
SEN-sEVs/SRT1720
E
Experimental schedule of drug administration
Cells xenografted
Chemo-administration
Tumor measurement
MIT
1st week
3rd week
5th week
7th week
Single or combined MIT/SRT2104 administration every other week
F
Transcript expression (Log2 fold change)
Placebo MIT Placebo MIT
Epithelial Stromal
IL6
IL8
IL1α
MMP3
WNT16B
GM-CSF
p16
SIRT1
G
Placebo
MIT
SRT2104
MIT/SRT2104
Vimentin/ABCB4
H
Log2 fold change
Placebo
MIT
SRT2104
MIT/SRT2104
Epithelial cells
Stromal cells

#### Supplementary Figure Legends

**Fig. S1 Cellular senescence, expression profiling and the SASP development in human stromal cells upon genotoxic treatment. Related to Fig. 1**

(A) Comparative statistics of SA-β-gal staining positivity in primary normal human prostate stromal cell line PSC27. Scale bars, 20 μm. (B) Comparative statistics of BrdU staining positivity in PSC27 cells. Scale bars, 10 μm. (C) DNA damage repair (DDR) profiling of PSC27 cells upon immunofluorescence staining for γ-H2AX. According to the number of DDR foci, cells were categorized into 4 subgroups: 0; 1-3; 4-10; >10. (D) Heatmap depiction of top 50 genes upregulated in SEN PSC27 cells after DNA damage treatment. (E) Comparative statistics of SA-β-gal staining positivity in primary human breast stromal cell line HBF1203. Scale bars, 20 μm. (F) Comparative statistics of BrdU staining positivity in HBF1203 cells. Scale bars, 10 μm. (G) Statistical DDR profiling of HBF1203 cells upon immunofluorescence staining for γ-H2AX, with a classification procedure described in C. (H) Transcript expression assay for HBF1203, including sEV biomarkers, sEV biogenesis-related molecules and the SASP hallmark factors measured. For each target, qRT-PCR signal of SEN cells was normalized to that of PRE cells. (I) Quantitative comparison of the number of sEVs secreted from PRE and SEN HBF1203 cells in 3 consecutive days as detected by NTA, data normalized to parental cell number. (J) NTA-based quantification of vesicles in P100 and P14 fractions after successive differential ultracentrifugation of conditioned media from PRE and SEN HBF1203 cells to measure the numbers of secreted sEVs and MVs, respectively. Data of A, B, C, E, F, G, H, I and J are representative of 3 independent experiments, with 3 technical replicates performed per experiment (^, $P > 0.05$; *, $P < 0.05$; **, $P < 0.01$; ***, $P < 0.001$).

**Fig. S2 Statistical analysis of miRNAs differentially carried by sEVs of PRE vs SEN stromal cells and database mapping of their targets in human. Related to Fig. 2**

(A) Venn diagram showing all miRNAs (both known and novel) detected by sRNA-Seq in sEVs derived from PRE and SEN stromal cells, with the total miRNA number per cell population and the number of miRNA overlapping between two populations presented. (B) Heatmap depicting the top upregulated (53) and downregulated (117) miRNAs carried in EVs from SEN vs PRE stromal cells. (C) Column presentation of the top 10 most abundantly expressed miRNAs (regardless of their identities are known or not) in each sequenced sample. (D) Column presentation of the top 10 most abundantly expressed miRNAs with known identities at miRBase in each sequenced sample. (E) Venn diagram showing the miRNA targets with available functional annotation presented by NR, Swiss-Prot, EggNOG, GO, KEGG and Pfam databases. (F) Gene Ontology (GO) analysis for top 50 molecular targets of the known miRNAs significantly upregulated in SEN vs PRE stromal cells with a fold change > 2.0 and $P < 0.05$. (G) Kyoto Encyclopedia of Genes and Genomes (KEGG) pathway analysis shows the functional involvement of the top 50 targets of known miRNAs significantly upregulated in SEN vs PRE stromal cells.

**Fig. S3 SIRT1 decline is not related with single strand breaks, proteasome degradation or autophagy deficiency upon cellular senescence. Related to Fig. 3**

(A) Quantitative measurement of HSP70, TSG101, IL8 and MMP3 transcript expression in stromal cells either naïve or upon DNA damage-induced senescence. (B) Immunoblot analysis of HSP70, TSG101, IL8 and MMP3 expression in PSC27 cells in conditions described in (A). GAPDH, loading control. (C) Left, immunoblots of HSP70, TSG101, ALIX and typical SASP markers (IL8 and MMP3) in stromal cells overexpressing exogenous human HSP70; right, NTA-based quantification of vesicles in P100 and P14 fractions post successive differential ultracentrifugation of stromal cell conditioned media. LE, long exposure, allowing signals to be visualized. GAPDH, loading control. (D) PSC27 cells were treated by Bay 11-7082 (BAY) and/or BLEO, and subject to immunoblot analysis 7 days afterwards. Cytoplasmic and nuclear fractions of a major subunit of the NF-κB complex, p65 (RELA), was assessed separately. GAPDH, loading control. (E) PSC27 cells were treated by Olaparib, a selective inhibitor of PARP1, and/or BLEO, with lysates of 7 days later subject to immunoblot analysis. (F) PSC27 cells were treated by either SAHA, a pan-HDAC inhibitor, or NAM, a SIRT1-specific inhibitor, and cell lysates were examined by immunoblots. LE, long exposure, allowing signals to be visualized. GAPDH, loading control. (G) Stromal cells were subject to BLEO and/or MG132, a potent cell-permeable proteasome inhibitor, and 7 days later lysates subject to immunoblot analysis. (H) PSC27 cells were treated by Bafilomycin A1, a vacuolar-ATPase inhibitor that disrupts late phase autophagy, and/or BLEO, with lysates of 7 days later assessed by immunoblots. (I) Stromal cells were treated by cycloheximide (CHX), an inhibitor of eukaryote protein synthesis, and/or BLEO, and subject to immunoblot examination 7 days later.

**Fig. S4 Senescent stromal sEVs enhance the malignancy of prostate cancer cells. Related to Fig. 4**

(A) Migration assay of PCa cells seeded within transwells in 6-well plates, after exposure to stromal sEVs for 3 days. Right, representative images of PC3 and DU145 migration measured via wound healing assay. Scale bar, 100 μm. (B) Invasiveness assessment of PCa cells across transwell membrane after exposure to stromal sEVs for 3 days. Right, representative images of PC3 and DU145 invasion across the transwell. Scale bar, 100 μm. (C) Dose-response curves (non-linear regression/curve fit) plotted from drug-based survival assays of PC3 cells exposed to sEVs collected from PRE or SEN HBF1203 cells, and subject to treatment by a wide range concentration of docetaxel (DOC). Whole

CM of stromal cells were used as experimental controls. (D) Apoptotic assay for combined activities of caspase 3/7 determined 24 hours after exposure of PC3 cells to sEVs of stromal cells while being treated by docetaxel (DOC) in the presence or absence of caspase inhibitors including QVD-OPH and ZVAD-FMK, or caspase activators including PAC1 and gambogic acid (GA). RLU, relative luciferase unit. Data of A, B, C and D are representative of 3 independent experiments, with 3 technical replicates performed per experiment (^, $P > 0.05$; *, $P < 0.05$; **, $P < 0.01$; ***, $P < 0.001$).

**Fig. S5 Transcriptomic profiling of prostate cancer cells upon exposure to senescent stromal sEVs and establishment of protein-protein interaction network involving ABCB4. Related to Fig. 5**

(A) Heatmap depicting the genes with significantly upregulated (221) or downregulated (123) expression in PC3 cells upon exposure to SEN stromal sEVs in relative to PRE stromal controls. (B) Scatter plot showing the significantly upregulated and downregulated genes of PC3 cells upon uptake of SEN stromal sEVs. (C) Heatmap depicting the significantly upregulated (191) and downregulated (231) genes in DU145 cells. (D) Scatter plot displaying the significantly upregulated and downregulated genes of DU145 cells after uptake of SEN stromal sEVs. In both (B) and (D), red, upregulated; green, downregulated; grey, insignificantly changed. (E) Heatmap showing the top 50 upregulated genes in PC3 cells after treatment with SEN stromal sEVs. (F) Heatmap showing the top 50 upregulated genes in DU145 cells after treatment with SEN stromal cell sEVs. (G) Gene Ontology (GO) analysis for the top 50 genes being significantly upregulated in PC3 cells after treatment with SEN stromal sEVs. (H) Gene Ontology (GO) analysis for the top 50 genes being significantly upregulated in DU145 cells after treatment with SEN stromal sEVs. (I) Venn diagram showing the number of genes being significantly upregulated by both PC3 and DU145 lines ($P < 0.05$ by *t* test, fold change > 2). (J) Heatmap displaying the 22 genes that were co-upregulated by PC3 and DU145 cells.

Red star, ABCB4. (K) Circos plot allowing visual presentation of genes correlated with the top 10 GO categories linked with the biological functions of 22 genes co-upregulated in two PCa cell lines upon exposure to SEN stromal sEVs. (L) Circos plot presenting correlation of genes with the top 4 KEGG categories relevant to the 22 co-upregulated genes in PCa cell lines upon exposure to SEN stromal sEVs. (M) Protein-protein interaction (PPI) network of protein molecules related to ABCB4. The nodes represent proteins, edges protein-protein associations.

**Fig. S6 Senescent stromal sEV-conferred malignancy is diminished upon elimination of ABCB4 from cancer cells. Related to Fig. 5**

(A) Immunoblot analysis of ABCB4 expression in PCa sublines. PC3 and DU145 cells were subject to ABCB4 knockdown mediated by lentiviral infection, followed by exposure to SEN vs PRE stromal sEVs. $shRNA^{C}$, scramble control. $shRNA^{ABCB4}$, ABCB4-specific shRNA (herein, #1 used as an example). GAPDH, loading control. (B) Migration assay of PCa cells exposed to SEN vs PRE stromal sEVs, with ABCB4 eliminated from these cells via shRNA-mediated knockdown. (C) Invasiveness examination of PCa cells treated by SEN vs PRE stromal sEVs, upon ABCB4 depletion from these cells. Data of (B) and (C) are representative of 3 independent experiments, with 3 technical replicates performed per experiment (^, $P > 0.05$; *, $P < 0.05$; **, $P < 0.01$; ***, $P < 0.001$).

**Fig. S7 Pharmacological targeting SIRT1 minimizes senescent stromal sEV production and prevents ABCB4 upregulation in prostate cancer cells. Related to Fig. 6**

(A) SIRT1 activation suppresses sEV production in SEN stromal cells. SRT1720, a selective SIRT agonist, was used to treat PSC27 cells at a concentration of 3 μM alone or together with BLEO at 50 μg/ml for 7-10 days, afterwards sEVs were collected in a 3-day time window and subject to NTA quantification. (B) Immunoblots of ABCB4 expression in PC3 cells upon incubation with sEVs from PRE or SEN stromal cells. GAPDH, loading control. (C) SIRT1 activation curbs sEV synthesis in SEN stromal cells of breast origin. SRT2104 or SRT1720 were used to treat HBF1203 breast stromal cells at 3 μM alone or together with doxorubicin (DOX) at 10 μM for 7-10 days, with collected sEVs subject to NTA measurement. (D) Dose-response curves plotted from drug-based survival assays of MDA-MB-231 breast cancer cells exposed to sEVs derived from HBF1203 treated by DOX and/or a SIRT1 agonist for 7-10 days, and concurrently exposed to a wide range of DOX concentrations. (E) Strategic workflow of drug administration and tumor surveillance for preclinical trial. Recombinant tissues composed of PC3/PSC27 cells ($1.25 \times 10^4$, in a ratio of 4:1) were inoculated subcutaneously to SCID mice 2 weeks before initiation of treatment. The chemotherapeutic agent MIT and the SIRT1 agonist SRT2104 were delivered simultaneously on the 1$^{st}$ day of each week starting from the 3$^{rd}$ week, then given every other week for a total number of 3 doses. At the end of the 8-week regimen mice were sacrificed with tumor volume measured, and histologically analyzed. (F) Transcript-based expression assay of several canonical SASP factors expressed in stromal cells isolated from the tumors of experimental mice. Tumor tissues from animals were subject to LCM isolation, RNA preparation and expression assessment. Signals were normalized to the lowest values among all examined samples. (G) Immunofluorescence staining of tumor tissues to locate the stromal and epithelial cells, and to examine ABCB expression level. Green, vimentin; red, ABCB4. Scale bar, 200 μm. (H) Measurement of ABCB4 expression in epithelial vs stromal cells dissected from mouse tumors and isolated with LCM. Signals were normalized to the lowest values among all examined samples. Data of A, C, D, F and H are representative of 3 independent experiments, with 3 technical replicates performed per cell-based assay (^, $P > 0.05$; *, $P < 0.05$; **, $P < 0.01$; ***, $P < 0.001$).

**Table S1. Statistics of small RNA species upon global assessment of Illumina sequencing data**

| Sample ID | PRE1 | PRE2 | PRE3 | SEN1 | SEN2 | SEN3 |
|---|---|---|---|---|---|---|
| Known miRNA | 2867 (0.06%) | 3109 (0.06%) | 2486 (0.06%) | 2718 (0.05%) | 2614 (0.05%) | 2749 (0.05%) |
| Novel miRNA | 140174 (3.04%) | 155585 (2.98%) | 152536 (3.51%) | 131801 (2.44%) | 115945 (2.22%) | 122040 (2.18%) |
| rRNA | 693254 (15.03%) | 820816 (15.74%) | 590941 (13.61%) | 505130 (9.36%) | 478052 (9.14%) | 535643 (9.55%) |
| tRNA | 354546 (7.69%) | 379560 (7.28%) | 280463 (6.46%) | 578650 (10.72%) | 652790 (12.48%) | 695134 (12.4%) |
| snoRNA | 0 (0.0%) | 0 (0.0%) | 0 (0.0%) | 0(0.0%) | 0 (0.0%) | 0 (0.0%) |
| snRNA | 3302 (0.07%) | 3724 (0.07%) | 2644 (0.06%) | 3568 (0.07%) | 3650 (0.07%) | 4177 (0.07%) |
| repbase | 28908 (0.63%) | 33653 (0.65%) | 28420 (0.65%) | 28573 (0.53%) | 28285 (0.54%) | 31476 (0.56%) |
| Exon | 22311 (0.48%) | 26351 (0.51%) | 20373 (0.47%) | 23834 (0.44%) | 21462 (0.41%) | 24693 (0.44%) |
| Intron | 134819 (2.92%) | 150428 (2.88%) | 123809 (2.85%) | 82971 (1.54%) | 79599 (1.52%) | 86608 (1.54%) |
| Unknown | 3233184 (70.08%) | 3643263 (69.84%) | 3141363 (72.33%) | 4041553 (74.86%) | 3846546 (73.56%) | 4105391 (73.21%) |
| Total | 4613365 | 5216489 | 4343035 | 5398798 | 5228943 | 5607911 |

**Table S2. Representative list of the molecular targets of upregulated and downregulated miRNAs in SEN stromal cell-derived sEVs**

| miRNA name | Target gene | Full name |
|---|---|---|
| ***Upregulated miRNAs*** | | |
| hsa-miR-6834-3p | THAP1 | THAP domain containing, apoptosis associated protein 1 |
| | NAIF1 | Nuclear apoptosis inducing factor 1 |
| | PERP | TP53 apoptosis effector |
| | DTHD1 | Death domain containing 1 |
| | PDCD6IP | Programmed cell death 6 interacting protein |
| | DAP | Death-associated protein |
| | ANKDD1A | Ankyrin repeat and death domain containing 1A |
| | PDCD1 | Programmed cell death 1 |
| | BID | BH3 interacting domain death agonist |
| hsa-miR-150-5p | AIFM2 | Apoptosis-inducing factor, mitochondrion-associated, 2 |
| | PDCD4 | Programmed cell death 4 (neoplastic transformation inhibitor |
| hsa-miR-136-3p | CASP7 | Caspase 7, apoptosis-related cysteine peptidase |
| | CASP8 | Caspase 8, apoptosis-related cysteine peptidase |
| | CASP10 | Caspase 10, apoptosis-related cysteine peptidase |
| | PDCD4 | Programmed cell death 4 (neoplastic transformation inhibitor |
| | PDCD5 | Programmed cell death 5 |
| | FAS | Fas cell surface death receptor |
| hsa-miR-542-3p | CDIP1 | Cell death-inducing p53 target 1 |
| | DEDD2 | Death effector domain containing 2 |
| hsa-miR-454-3p | BCL2L11 | BCL2-like 11 (apoptosis facilitator) |
| | TNF | Tumor necrosis factor |
| | DEDD | Death effector domain containing |
| ***Downregulated miRNAs*** | | |
| hsa-miR-130b-3p | ABCB7 | ATP-binding cassette, sub-family B (MDR/TAP), member 7 |
| hsa-miR-214-3p | ABCB1 | ATP-binding cassette, sub-family B (MDR/TAP), member 1 |
| | ABCB8 | ATP-binding cassette, sub-family B |

| | | |
|---|---|---|
| | | (MDR/TAP), member 8 |
| | ABCB9 | ATP-binding cassette, sub-family B (MDR/TAP), member 9 |
| | ABCB10 | ATP-binding cassette, sub-family B (MDR/TAP), member 10 |
| hsa-miR-7704 | ABCB8 | ABCB8 ATP-binding cassette, sub-family B (MDR/TAP), member 8 |
| | API5 | Apoptosis inhibitor 5 |
| | XIAP | X-linked inhibitor of apoptosis |

**Table S3. Predicted miRNA target molecules located at the key nodes of intracellular protein-protein interaction (PPI) network**

| Node1 | Node2 | node1_accession_id | node2_accession_id |
|---|---|---|---|
| **TNFRSF8** | PAX5 | ENSG00000120949 | ENSG00000196092 |
| **CD72** | PAX5 | ENSG00000137101 | ENSG00000196092 |
| **TOMM70A** | FKBP5 | ENSG00000154174 | ENSG00000096060 |
| **CD72** | MMP14 | ENSG00000137101 | ENSG00000157227 |
| **MSI1** | UBFD1 | ENSG00000135097 | ENSG00000103353 |
| **UBE2R2** | UBFD1 | ENSG00000107341 | ENSG00000103353 |
| **UBE2R2** | FKBP5 | ENSG00000107341 | ENSG00000096060 |

#### Table S4. Functional enrichments of predicted partners of ABCB4 in PPI network

##### Biological Process (GO)

| *Description* | *Count in geneset* | *False discovery rate* |
|---|---|---|
| Regulation of lipid metabolic process | 11 of 373 | 1.32E-16 |
| Positive regulation of transcription by RNA polymerase II | 8 of 1104 | 6.91E-06 |
| Positive regulation of RNA metabolic process | 8 of 1596 | 4.16E-05 |
| Steroid hormone mediated signaling pathway | 4 of 131 | 6.64E-05 |
| Monocarboxylic acid transport | 4 of 131 | 6.64E-05 |

##### Molecular Function (GO)

| *Description* | *Count in geneset* | *False discovery rate* |
|---|---|---|
| Transcription coregulator activity | 7 of 534 | 5.82E-07 |
| Transcription regulatory region DNA binding | 7 of 829 | 5.88E-06 |
| Transcription coactivator activity | 5 of 307 | 1.40E-05 |
| DNA binding | 9 of 2457 | 1.40E-05 |
| Transcription regulator activity | 8 of 2069 | 5.32E-05 |

##### Cellular Component (GO)

| *Description* | *Count in geneset* | *False discovery rate* |
|---|---|---|
| Nucleoplasm | 11 of 3446 | 4.93E-07 |
| Spindle microtubule | 2 of 47 | 0.0032 |
| Transcriptional repressor complex | 2 of 80 | 0.0081 |
| Chromatin | 3 of 489 | 0.0165 |
| RNA polymerase II transcription factor complex | 2 of 139 | 0.0174 |

##### KEGG Pathways

| *Description* | *Count in geneset* | *False discovery rate* |
|---|---|---|
| PPAR signaling pathway | 3 of 72 | 0.00037 |
| Thyroid hormone signaling pathway | 3 of 115 | 0.00073 |
| Wnt signaling pathway | 3 of 143 | 0.00092 |
| Bile secretion | 2 of 71 | 0.0075 |
| Adipocytokine signaling pathway | 2 of 69 | 0.0075 |

##### Reactome Pathways

| *Description* | *Count in geneset* | *False discovery rate* |
|---|---|---|
| RORA activates gene expression | 9 of 18 | 2.50E-23 |
| PPARA activates gene expression | 11 of 114 | 2.59E-23 |
| BMAL1:CLOCK,NPAS2 activates circadian gene expressi | 9 of 27 | 1.26E-22 |
| Activation of gene expression by SREBF (SREBP) | 9 of 40 | 2.19E-21 |
| Transcriptional activation of mitochondrial biogenesis | 9 of 53 | 1.80E-20 |

##### UniProt Keywords

| *Description* | *Count in geneset* | *False discovery rate* |
|---|---|---|
| Transcription regulation | 9 of 2279 | 8.67E-06 |
| Activator | 6 of 670 | 1.08E-05 |
| Nucleus | 9 of 5200 | 0.0026 |
| Chromatin regulator | 3 of 287 | 0.0048 |
| Acetylation | 7 of 3335 | 0.006 |

##### PFAM Protein Domains

| *Description* | *Count in geneset* | *False discovery rate* |
|---|---|---|
| LisH | 2 of 6 | 0.00026 |
| Anaphase-promoting complex subunit 4 WD40 domain | 2 of 30 | 0.0023 |
| Zinc finger, C4 type (two domains) | 2 of 46 | 0.0035 |
| Ligand-binding domain of nuclear hormone receptor | 2 of 48 | 0.0035 |

| | | |
|---|---|---|
| WD domain, G-beta repeat | 2 of 224 | 0.045 |

**INTERPRO Protein Domains and Features**

| ***Description*** | ***Count in geneset*** | ***False discovery rate*** |
|---|---|---|
| LIS1 homology motif | 2 of 27 | 0.0067 |
| Nuclear hormone receptor-like domain superfamily | 2 of 48 | 0.0093 |
| G-protein beta WD-40 repeat | 2 of 85 | 0.0093 |
| Zinc finger, NHR/GATA-type | 2 of 56 | 0.0093 |
| Nuclear hormone receptor | 2 of 46 | 0.0093 |

#### Table S5. Primer sequences for qRT-PCR assays

| Gene name | Forward (5’-3’) | Reverse (5’-3’) |
|---|---|---|
| ***IL-6*** | TACCCCCAGGAGAAGATTCC | TTTTCTGCCAGTGCCTCTTT |
| ***IL-8*** | GTGCAGTTTTGCCAAGGAGT | CTCTGCACCCAGTTTTCCTT |
| ***IL-1α*** | AATGACGCCCTCAATCAAAG | TGGGTATCTCAGGCATCTCC |
| ***WNT16B*** | GCTCCTGTGCTGTGAAAACA | TGCATTCTCTGCCTTGTGTC |
| ***MMP3*** | GCAGTTTGCTCAGCCTATCC | GAGTGTCGGAGTCCAGCTTC |
| ***GM-CSF*** | ATGTGAATGCCATCCAGGAG | AGGGCAGTGCTGCTTGTAGT |
| $p16^{INK4a}$ | CTTCCTGGACACGCTGGT | GCATGGTTACTGCCTCTGGT |
| ***SIRT1*** | GCAGATTAGTAGGCGGCTTG | TCTGGCATGTCCCACTATCA |
| ***ATP6V1A*** | ATGACAGGTTCTGCCCATTC | GAGGATGTCTCCCATGTGCT |
| ***ABCB4*** | GCATCAGCAGCAAACAAAAA | GCAGCGACAAGGAAAAGTTC |
| ***RPL13A*** | GTACGCTGTGAAGGCATCAA | CGCTTTTTCTTGTCGTAGGG |
| ***TSG101*** | GATACCCTCCCAATCCCAGT | GTCACTGACCGCAGAGATGA |
| ***Syntenin-1*** | TTCTGCTCCTATCCCTCACG | CCAGTTACAGGAGCCACCAT |
| ***ALIX*** | TGGCTGCAAAGCACTGTATC | AGGGCACGATTGATTTTGTC |
| ***CD9*** | TTGGACTATGGCTCCGATTC | GGCGAATATCACCAAGAGGA |
| ***CD63*** | TCCCTTCCATGTCGAAGAAC | TCCCAAAACCTCGACAAAAG |
| ***CD81*** | CTGCACCAAGTGCATCAAGT | CCAACGAACATCATGACAGC |
| ***ADAM10*** | AGCAACATCTGGGGACAAAC | CCCAGGTTTCAGTTTGCATT |
| ***EHD4*** | AAAGACAAGCCCGTCTACGA | CGAACTCCTCCTCATCAAGC |
| ***SMPD2*** | TTTGGTGTCCGCATTGACTA | TAGAGCTGGGGTTCTGCTGT |
| ***SMPD3*** | AAGAGGACAGCCTCTGTGGA | CTGATTATGGTTGGGCGTCT |

**Table S6. Antibodies used for immunoblot, immunofluorescence and immunohistochemistry staining**

| Antigen name | Commercial source | Catalog number (clone number) | Application |
|---|---|---|---|
| **HSC70** | Santa Cruz Biotechnology | sc-7298(B-6) | Western blot |
| **Syntenin-1** | Abcam | ab133267(EPR8102) | Western blot |
| **TSG101** | Santa Cruz Biotechnology | sc-7964(C-2) | Western blot, ELISA,IF |
| **ALIX** | Cell Signaling Technology | 2171(3A9) | Western blot |
| **CD63** | Bioss | bs-1523R(4C2) | Western blot, IF |
| **CD81** | Abnova | PAB16754 | Western blot |
| **BrdU** | Cell signaling | 5292 | IF |
| **SIRT1** | Bimake | A5049 | Western blot, IHC |
| **SIRT2** | abcam | ab67299 | Western blot |
| **ATP6V1A** | Proteintech | 17115-1-AP | Western blot |
| **p62** | Bimake | A5180 | Western blot |
| **LC3II** | Bimake | A5202 | Western blot |
| **IL8** | Abcam | AB18672(807) | Western blot |
| **MMP3** | Proteintech | 66338-1-Ig | Western blot |
| **EGFR** | Santa Cruz Biotechnology | sc-03-G(A-10) | Western blot |
| **Ubiquitin** | Abway | F040501(CY5520) | Western blot, IHC |
| **Caspase 3 (cleaved)** | Cell signaling | 9661 | WB, IHC |
| **Caspase 3 (intact)** | Cell signaling | 9662 | WB |
| **$p16^{INK4a}$** | GeneTex | GTX115342 | IHC |
| **ABCB4** | Fitzgerald Industries | 70R-6711 | IHC |
| **GAPDH** | Abway | F040501(AB0037) | Western blot |
